# Biomolecular condensates sustain pH gradients at equilibrium through charge neutralisation

**DOI:** 10.1101/2024.05.23.595321

**Authors:** Hannes Ausserwöger, Rob Scrutton, Charlotte M. Fischer, Tomas Sneideris, Daoyuan Qian, Ella de Csilléry, Ieva Baronaite, Kadi L. Saar, Alan Z. Białek, Marc Oeller, Georg Krainer, Titus M. Franzmann, Sina Wittmann, Juan M. Iglesias-Artola, Gaetano Invernizzi, Anthony A. Hyman, Simon Alberti, Nikolai Lorenzen, Tuomas P. J. Knowles

## Abstract

Electrochemical gradients are essential to the functioning of cells and form across membranes using active transporters. Here, we show in contrast that condensed biomolecular systems sustain significant pH gradients without any external energy input. By studying individual condensates on the micron scale using a microdroplet platform, we reveal dense phase pH shifts towards conditions of minimal electrostatic repulsion. We demonstrate that by doing so protein condensates can drive substantial alkaline and acidic gradients which are compositionally tuneable and can extend to complex architectures sustaining multiple unique pH conditions simultaneously. Through in silico characterisation of human proteomic condensate networks, we further highlight potential wide ranging electrochemical properties emerging from condensation in nature, while correlating intracellular condensate pH gradients with complex biomolecular composition. Together, the emergent nature of condensation shapes distinct pH microenvironments, thereby creating a unique regulatory mechanism to modulate biochemical activity in living systems.

## Introduction

The proton (H^+^) concentration, typically represented as the pH, is a uniquely important environmental factor in biology. It affects all fundamental processes underlying viability and functioning of cells, including electrochemical potential gradients^1^, chemical reaction rates^2^ or macromolecular conformation and assembly^3,4^. pH mediated changes in the functional behaviour of proteins, for example, are driven by the inherent ability of functional groups to display different protonation states, shifting their physico-chemical properties. As a consequence, pH is tightly regulated within the intracellular environment, with deviations from the cytosolic pH only common in membrane-bound organelles^5,6^. Through energy input using active transporters, the organellar pH is regulated to optimise the specific reactions and processes performed in the organelle^7–12^. Precise control over the intracellular pH is further aided by a high buffering capacity of the cytosol, which originates primarily from smaller ionic species such as phosphates and carbonates^5,6^. In comparison, the concentration of protein associated ionisable side groups is typically far too low to contribute, accounting to only about 1% of the cytosolic buffering capacity in yeast cells^13^.

Remarkably, this low-protein-concentration picture can be drastically altered as a result of biomolecular phase separation, leading to the formation of a protein-rich dense phase and a protein-poor dilute phase. Here, collective interactions drive local concentration increases of biomolecules in the condensed state to give rise to unique chemical microenvironments at equilibrium^14,15^. These unique microenvironments, often referred to as condensates, are believed to play critical roles in cellular organisation and pathology^14,16,17^. They can yield distinct material properties or differential molecular partitioning^18^, which has even been shown to extend to the formation of elemental ion gradients^19,20^. Reports even suggest that condensates could drive pH gradients intracellularly^21–26^. Hence, we sought a generalisable understanding of how pH influences and drives the collective assembly of biomolecules as well as how the internal environment of these assemblies can modulate external pH conditions and establish pH gradients.

## Results

### Mapping pH-responsive phase boundaries in a continuous manner

To quantify pH-responsive phase behaviour, we employed a combinatorial droplet microfluidic platform, enabling the generation of large numbers of water-in-oil droplets with distinct chemical microenvironments (Fig. 1a, b)^27^. Continuous variation of the pH in droplets was achieved by integrating a histidine and succinic acid buffer system (H/S buffer), capable of regulating pH between 3.5 – 9, while displaying a linear correlation between the pH and the volume fraction of two separate buffer stocks (see Materials and Methods and SI Fig. 1). These individual aqueous droplets were then subjected to image analysis to characterise the polypeptide phase state in each distinct chemical microenvironment. We first selected insulin and its clinically relevant variant, insulin glargine, because of their well-characterized pH-sensitive self-assembly^28–30^ and availability as model systems, enabling a direct assessment of how subtle sequence modifications influence phase separation across varying pH conditions. Application of this platform to insulin glargine (modified insulin variant, see SI Table 1 for sequences) yielded a continuous map of over 75000 data points with varying pH and insulin glargine concentrations, with each providing information on the local phase behaviour (Fig. 1c, SI Fig. 2 and 3 for data processing).

**Figure 1:**
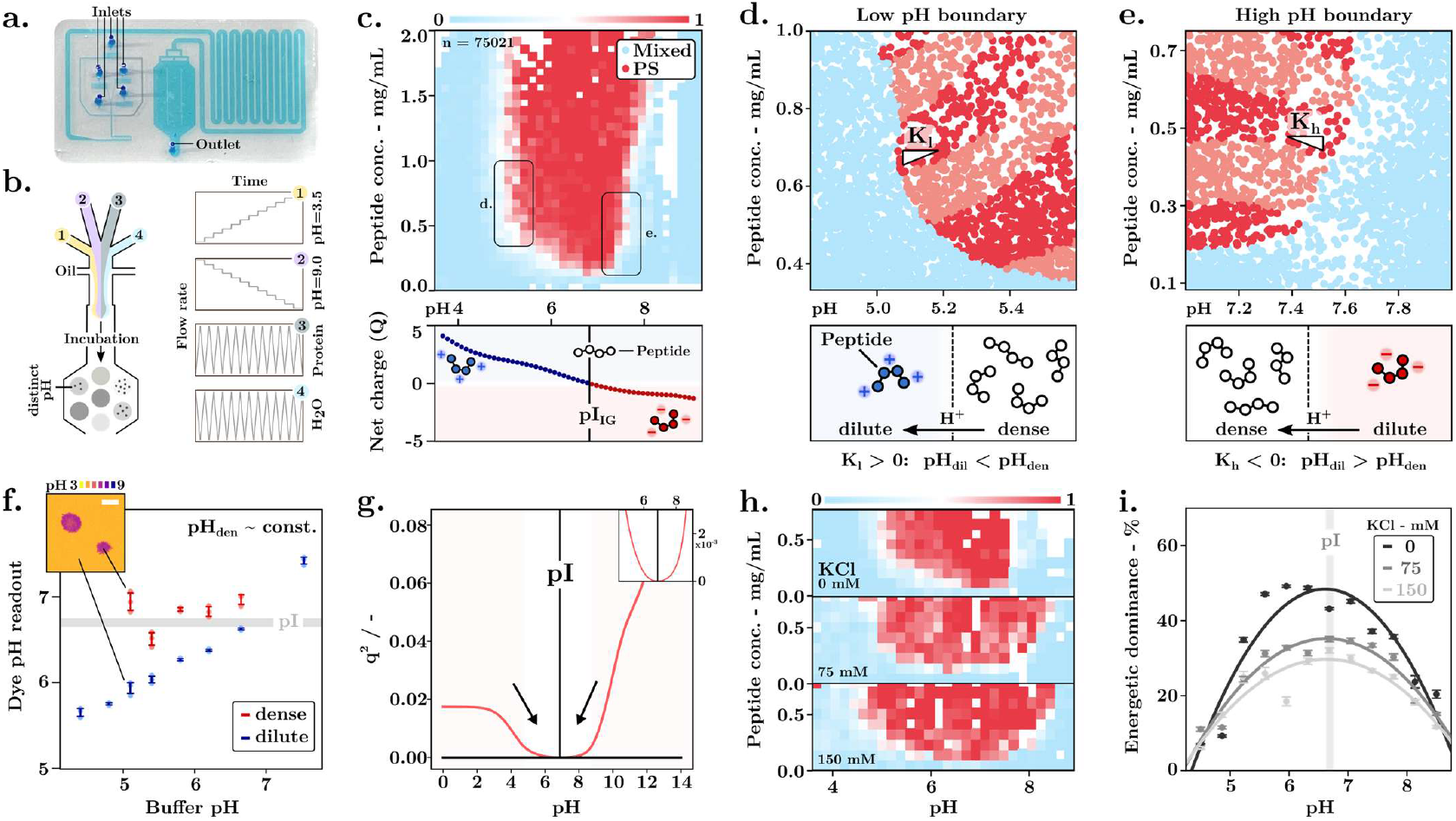
Deciphering pH-responsive macromolecular phase behaviour. **(a)** Image of the microfluidic chip, filled with a coloured solution to highlight the device features, used to generate continuous pH phase diagrams. **(b)** Schematic of the chip design and flow profiles used to generate continuous pH phase boundaries. **(c)** pH-responsive phase boundary of an insulin variant peptide (insulin glargine), correlated with the molecular net charge (Q). Data are binned into a grid-map to evaluate the local phase separation probability (P_PS_ = 1: red, P_PS_ = 0.5: white, P_PS_ = 0: blue) as determined by averaging (n_tot_ = 75021, see Supporting Information). **(d, e)** Visualisation of the dilute phase contour gradients (K) at the low pH (d) and high pH (e) phase boundary. Dilute phase contours are displayed using red shading of phase separated points as generated by sectioning for the dilute phase concentration obtained in each individual microdroplet^19,32,34^. Lower panels: Schematic representations visualising the proton partitioning as inferred from the reduced tie line gradients. **(f)** Dense and dilute phase pH readouts of insulin glargine samples at varying pH conditions using the ratiometric pH sensing dye SNARF-4F. Inset: representative image colour-coded for the pH readout. Scale bar: 10 μm. Data are presented as mean values ± SD. **(g)** Change in the square of the sequence charge density (q) used as a proxy for the repulsion free energy with varying pH for insulin glargine. **(h)** Effect of the addition of KCl on the pH-responsive phase boundary of the insulin glargine (n_75mM_ = 12623, n_150mM_ = 9969). **(i)** Quantification of the protein dominance with varying pH and salt concentrations for the insulin glargine. Data are presented as mean values ± SD.

Phase separation of insulin glargine was only observed close to the isoelectric point (pI) at 6.8, highlighting the importance of the sequence derived net charge (Q) profile. This suggests that molecular charge neutrality is a requirement for the intermolecular interactions to become favourable. Insulin (pI_Ins_ = 5.4; glargine variant: N21G mutation, 2x R additions in the β-chain) displays a similar correlation between the phase boundary and the sequence charge, where in both cases the minimum in the critical concentration coincides with the pI (see SI Fig. 4). Accordingly, the dilute phase concentration (c_dil_) also displayed a minimum around the pI, meaning that phase separation propensity attains a maximum at the pI (see SI Fig. 5a).

A decrease in the width of the pH phase boundary of insulin glargine compared to unmodified insulin (see SI Fig. 4) coincides with an increase in the overall sequence hydrophilicity (H_pH=7, I_ = 14, H_pH=7, IG_ = 21; Kyte-Doolittle scale^31^). Addition of two charged arginine residues makes insulin less hydrophilic, indicating a trade-off between intermolecular charge repulsion and attractive Van der Waals interactions. It is worth noting that the phase boundary is not perfectly symmetrical suggesting higher order effects form the individual ionisation changes of specific amino acids.

### Polypeptide chains buffer pH towards charge-neutral conditions

Next, we set out to characterise the proton partitioning between the dense and dilute phase by applying a previously published partitioning analysis approach^19,32,33^. Briefly, we quantified the peptide dilute phase concentrations in each individual water-in-oil droplet and investigated its change under varying pH and peptide concentration conditions. In an exact two-component system, contours of constant c_dil_ coincide with tie lines connecting the dilute and dense phase composition^19,34^ (see SI section ‘Reduced tie line approach in pH chemical space’). Visualisation of these contours is achieved by sectioning the dilute phase concentration conditions in set thresholds of a < c_dil_ < b (see Fig. 1d, e; light and dark red shaded contours, see SI Fig. 2 for data processing). Other species, however, affect the slope of mentioned contours, referred to here as K, therefore, only providing a lower bound for tie line component ratio^32^. This distortion is dependent on the shape of the phase boundary, with minimal effect where the phase boundary is parallel to the peptide axis such as at the extreme pH conditions of the observed phase boundary^32^.

When triggering insulin glargine phase separation at low pH (~ 5), the dilute phase contour shows a positive gradient (see Fig. 1d, K_l_ > 0), indicating that the pH in the dilute (pH_dil_) and dense phase (pH_den_) are not equivalent. Specifically, this suggests that pH_den_ > pH_dil_, i.e. protons are being excluded from the dense phase to bring the dense phase pH closer to the pI at 6.8. At the high pH phase boundary (around pH ~ 8) we find differential partitioning of protons into the dense phase (K_h_ < 0) to again bring the dense phase pH closer to the pI compared to the dilute phase (see Fig. 1e). Hence, independently of the starting pH, this would yield a decrease in the molecular net charge of the peptide in the dense phase and allow for a reduction of intermolecular repulsion. A similar behaviour is observed for insulin at the high pH boundary where K_pH_ < 0 (see, SI Fig. 5b). The low pH phase boundary could not be resolved with the available buffer system. We next set out to orthogonally confirm the observation of pH gradients between the dense and dilute phase. We applied the ratiometric pH sensing dye SNARF-4F and read out the dye signal in a spatially resolved manner using confocal imaging (see SI Fig. 6). Using this approach, we determine dilute and dense phase pH of insulin glargine samples at varying initial pH conditions (see Fig. 1f). While the dilute phase pH changes as expected with the starting pH, the apparent dense phase pH was observed to stay constant between ~ 6.5 and 7. This confirms the regulation of the dense phase pH towards the polypeptide chain isoelectric point (pI = 6.8) and supports the hypothesis of driving pH gradients to decrease electrostatic repulsion. Indeed, unmodified insulin samples display a similar trend where the apparent dense phase pH is constant while the dilute phase pH varies with the starting pH condition (see SI Fig. 5c). As such, the condensed phase effectively appears to act as a spatially localised buffer that can shift the pH conditions to minimise charge repulsion. To quantify the potential origin of the dense phase buffering effect, we measured the insulin c_dil_ at varying concentrations of H/S buffer but fixed pH = 6.4 and total peptide concentration (SI Fig. 7). Here, c_dil_ decreases with decreasing H/S buffer concentration highlighting a competition between the environmental buffer strength and phase separation. This can be explained as the dense phase has to act as a distinct buffering system against the environmental pH to generate the pH gradient necessary to minimise repulsion. By combining data sets at 2 and 1.5 mg/mL total insulin concentration, we can then also perform partitioning analysis for the H/S buffer^19,34^. We find a negative reduced tie line gradient (SI Fig. 7), i.e. that the buffer concentration is lower in the dense phase than the dilute phase. Hence, regulation of the dense phase pH is aided by buffer exclusion to favour the buffering of protein associated side groups. Such protein buffering is expected given common dense phase concentration in the low mM range^35–37^, meaning that the side-chain concentration will be comparable to or even exceed the environmental buffering.

### pH gradients reduce intermolecular electrostatic repulsion of condensation

We next set out to study change in the dense phase repulsion free energy with pH. Using a scaling argument, we suggest that the dense phase repulsion free energy scales with the square of sequence charge density (q) of the polypeptide chain, as the repulsion free energy of assembling a charged sphere of fixed radius is proportional to the square of its charge density^38^. While the dense phase concentration (c_den_), among other factors, will modulate the repulsion free energy, q^2^ can serve as a proxy for the relative energetic change. The sequence charge density was then calculated by normalising the peptide net charge at varying pH, as obtained from tabled pK_a_ values of the individual amino acids, by the sequence length N (q = Q/N). The q^2^ profile for insulin glargine displays a minimum around the pI, with a sharp increase at around pH 5 and 8 (Fig. 1g), corroborating that the observed pH gradients help mitigate electrostatic repulsion in the dense phase. The repulsion profile also resembles the overall phase boundary, which is similarly observed for human insulin (SI Fig. 5d).

To test this further we sought to modulate the electrostatic repulsion through addition of potassium chloride (KCl) and associated charge screening. At fixed KCl concentrations of 75 and 150 mM, an increase in the width of the pH-responsive phase boundary for insulin glargine was observed (see Fig. 1h). Hence, phase separation could occur at higher polypeptide net charge due to the added charge shielding. Differential shifts of the boundary in the high pH regime compared to the low pH regime are likely associated with ion-side chain specific interactions or potential charge regulation effects aided by the presence of ions.

To quantify the effect of the counter ions on the energetics, we next applied a previously published dominance analysis approach^19,32,33^. This allows us to map the relative free energy gain (D_i_) of individual components to the overall energetic driving force of phase separation. To do so we follow trajectories of increasing insulin glargine concentrations at fixed pH in the microdroplet pH scan (SI Fig. 8). The dilute phase response gradient R_AA_ at insulin glargine concentrations above c_sat_ can then be evaluated to calculate the dominance (D = 1 – R). At a pH of 6.1 and in the absence of salt, for example, we observe a D_p_ = 0.46 for insulin glargine, showing that even under salt-free conditions other factors contribute to the free energy gain associated with phase separation. We also find that D_p_ is maximised at pHs around the peptide pI (Fig. 1i), highlighting that the protein behaves much more like a one component system at low peptide net charge. Upon addition of KCl this trend is conserved but we observe a general decrease in the insulin glargine dominance with increasing the KCl concentrations, as ions are starting to increasingly contribute to energetic driving force through their electrostatic modulation effects.

### PGL3 condensation creates acidic pH microenvironments

We reason that the necessity to reduce electrostatic repulsion via pH gradients in the dense phase will also translate to other protein systems and conditions. To test this, we investigated the condensation of PGL3, a protein critical to the germ cell development of *Caenorhabditis elegans* by scaffolding the formation of P-granules^39^. Instead of varying the pH condition we induced PGL3 phase separation by decreasing the ionic strength at fixed pH (50 mM TRIS, pH = 7.3) in presence of SNARF-4F. We find an acidification of the dense phase compared to the dilute phase (Fig. 2a, b and c), which was observed to be independent of condensate size (SI Fig. 9). The resulting acidic dense phase pH gradient brings the dense phase pH significantly closer to the isoelectric point of the protein (pI_PGL3_ = 5.1 from sequence-based prediction; pI value only minimally effected by folding changes, see SI Table 2). In agreement with previous observations the dense phase pH shift decreases intermolecular PGL3 repulsion, as confirmed by a minimum in the q^2^ profile at acidic pH (see Fig. 2d). The difference between the apparent dense phase pH at around 6-6.5 compared to the pI can be explained by the asymmetry in the q^2^ profile of PGL3 towards a smaller repulsion increase at pH > pI together with the presence of electrostatic shielding of the ions present.

**Figure 2:**
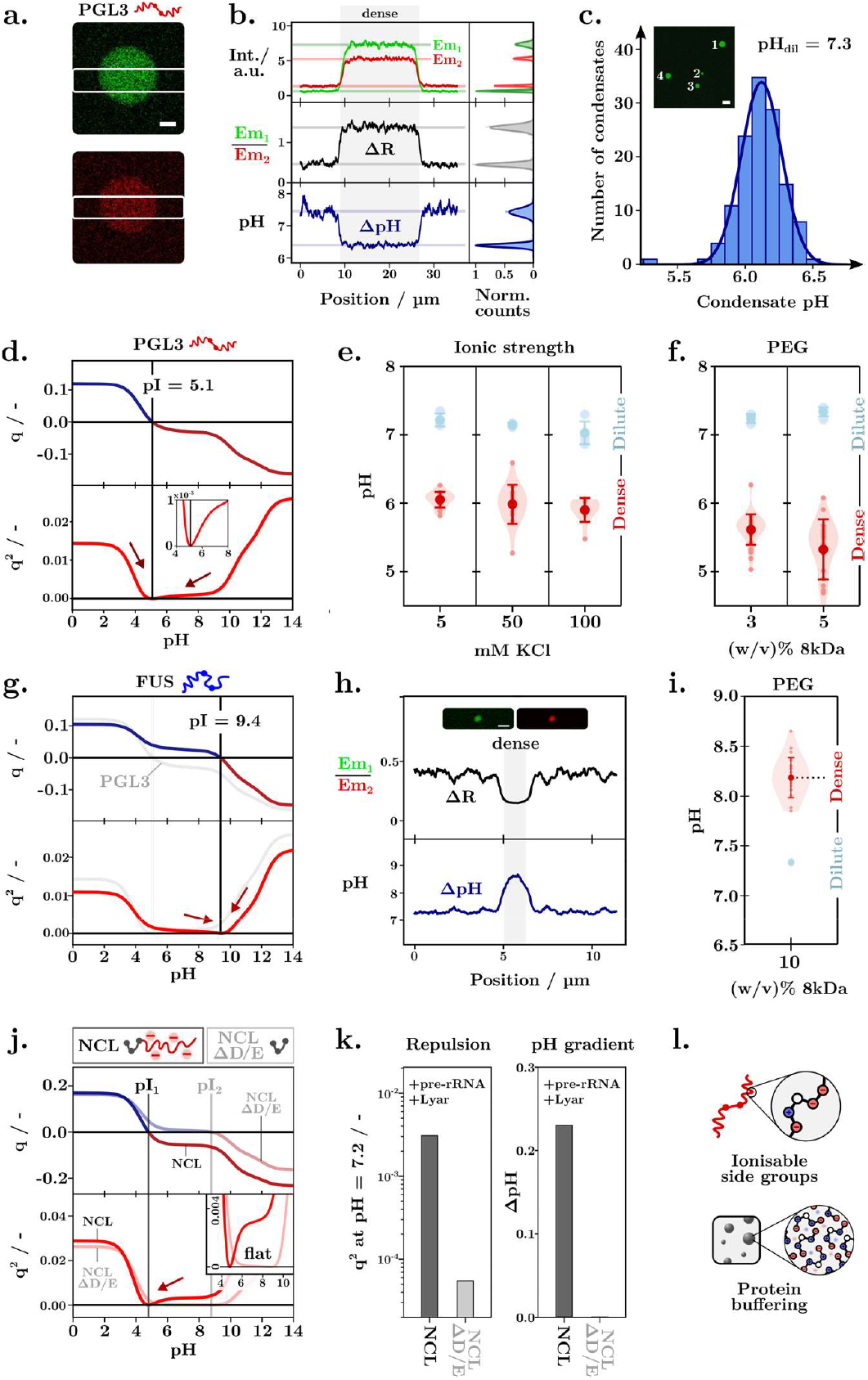
Biomolecular condensates sustain dense phase pH gradients buffered towards the protein pI. **(a)** Confocal imaging of PGL3 condensates at 2 μM protein concentration and 1 mM KCl, 50 mM TRIS buffer at pH = 7.3 using the ratiometric pH sensing dye SNARF-4F. Scale bar: 5 μm. **(b)** Spatially resolved ratiometric dye signal in dense and dilute phase for both emission channels, the emission intensity ratio and the corresponding pH, including readout histograms of all signal traces. **(c)** Condensate pH distribution for large numbers of PGL3 condensates formed at low ionic strength conditions from (a). The dilute phase pH was measured at 7.3. Inset: Confocal image of multiple condensates. Scale bar: 20 μm. **(d)** Net charge density and net charge density square of PGL3 protein with varying pH. **(e, f)** PGL3 dense and dilute phase pH at varying KCl and PEG concentrations. Fixed total protein concentration and buffer condition (2 μM protein, pH=7.3, 50 mM TRIS buffer, 150mM KCl in g and h). Data are presented as mean values ± SD. **(g)** Net charge density and net charge density square of FUS protein with varying pH. **(h)** Spatially resolved ratiometric dye signal in dense and dilute phase for both emission channels, the emission intensity ratio and the corresponding pH, for a FUS condensate formed at 10 (w/v)% PEG, 2.5 μM protein, 150 mM KCl and 50 mM TRIS buffered at pH=7.3. Scale bar: 2 μm **(i)** FUS dense and dilute phase pH at 10 (w/v)% PEG, 2.5 μM protein, 150 mM KCl and 50 mM TRIS buffered at pH=7.3. Data are presented as mean values ± SD. **(j)** Comparison net charge density and net charge density square for the NCL protein and ΔD/E NCl deletion mutation (deletion of a sequence region rich in acidic amino acids – D/E tract). **(k)** Comparison of the net charge density square at pH = 7.2 for NCL with and without the D/E tract (left). Observed *in vitro* pH gradient from ^26^ for condensates formed with 15 μM NCL protein, 1 μM pre-rRNA and 0.2 μM Lyar protein. **(l)** Schematic representation of the mechanism of action underlying the protein dense phase buffering driving pH gradients.

We next subjected PGL3 to phase separation under varying KCl and polyethylene glycol (PEG) concentrations (Fig. 2e, f), to understand the impact of different solute constitutions on the observed pH gradient. Changes in the KCl concentration between 5-100 mM did not change pH_den_ significantly as all conditions display an apparent dense phase pH < 6.5. Hence, for PGL3, independent of the environmental ionic strength, the formation of a dense phase pH gradient is still necessary to reduce protein-protein repulsion. When triggering PGL3 phase separation by introducing PEG we also find an acidic dense phase pH closer to the protein pI. This shows that the formation of a PGL3 condensed phase requires sustaining a pH gradient even at higher ionic strengths and independently of the phase separation trigger.

### Effective repulsion governs condensate pH gradients

Accordingly, highly positively charged proteins could yield the formation of alkaline condensate pH gradients. To test this, we turned to the FUS protein with pI = 9.4 (see Fig. 2g) which has been associated with aberrant phase transitions in the emergence of amyotrophic lateral sclerosis (ALS)^40^. Indeed, when triggering FUS phase separation by addition of PEG we find an apparent dense phase pH > 8 using SNARF-4F readouts (see Fig. 2h, i). Similarly to PGL3, FUS sets up a pH gradient towards its pI under a range of other homotypic phase separation conditions (see SI Fig. 10), highlighting that condensates can sustain pH buffering towards a variety of pH conditions. The condensed-phase buffering effect is corroborated by an observed competition between FUS phase separation and the environmental buffer (SI Fig. 11). Notably, lowering the buffer concentration can induce condensation. Consequently, decreasing buffer concentrations reduces the energetic barrier for the condensed phase to regulate pH_den_, thus diminishing repulsive interactions and promoting phase separation.

To investigate how the effective protein-protein repulsion modulates the observed condensate pH gradient we turned to previously published work on a NCL protein system associated with nucleolar phase separation^26^ (see Fig. 2j). The negatively charged wildtype NCL protein (pI = 4.8) forms a slightly acidic dense phase when undergoing phase separation with RNA, in presence of low concentrations of another client protein (LYAR, 70-fold lower concentration). This is expected when evaluating the q^2^ profile of NCL which highlights a clear minimum at low pH (Fig. 2k). Deletion of a region rich in negatively charged amino acids from the protein (ΔD/E NCl, pI = 8.8) also removes the presence of a dense pH gradient. While the ΔD/E NCl variant displays a high pI it effectively presents zero net charge between pH 5-9 (Fig. 2j). This yields minimal repulsion in the dense phase even at pH conditions different from the pI, suggesting that sustaining pH gradients only becomes necessary at sufficient levels of electrostatic repulsion at the environmental pH. Together, these data highlight that biomolecular condensates can regulate the dense phase pH towards the pI through the proteins innate buffering capacity under environmental conditions that are electrostatically unfavourable for phase separation (Fig. 2l).

### Effect of RNA-scaffolding on condensate pH gradients

Protein phase separation commonly arises from heterotypic interactions especially with nucleic acids, leading us to further investigate the effect of heteromolecular mixtures on the condensate pH. When triggering PGL3 phase separation by addition of polyadenylic acid RNA (p(A)-RNA), we find an acidic apparent pH_den_ of around 6 (Fig. 3a) comparable to the homotypic PGL3 condensates (see Fig. 2c, e). Crucially, nucleic acids are also characterised by a low pI because of their phosphate backbone groups. Therefore, heterotypic PGL3-RNA interactions will also become more favourable at lower pH because of a decrease in repulsion. The co-partitioned RNA will also contribute to decreasing the dense phase pH via buffering of its phosphate backbones.

**Figure 3:**
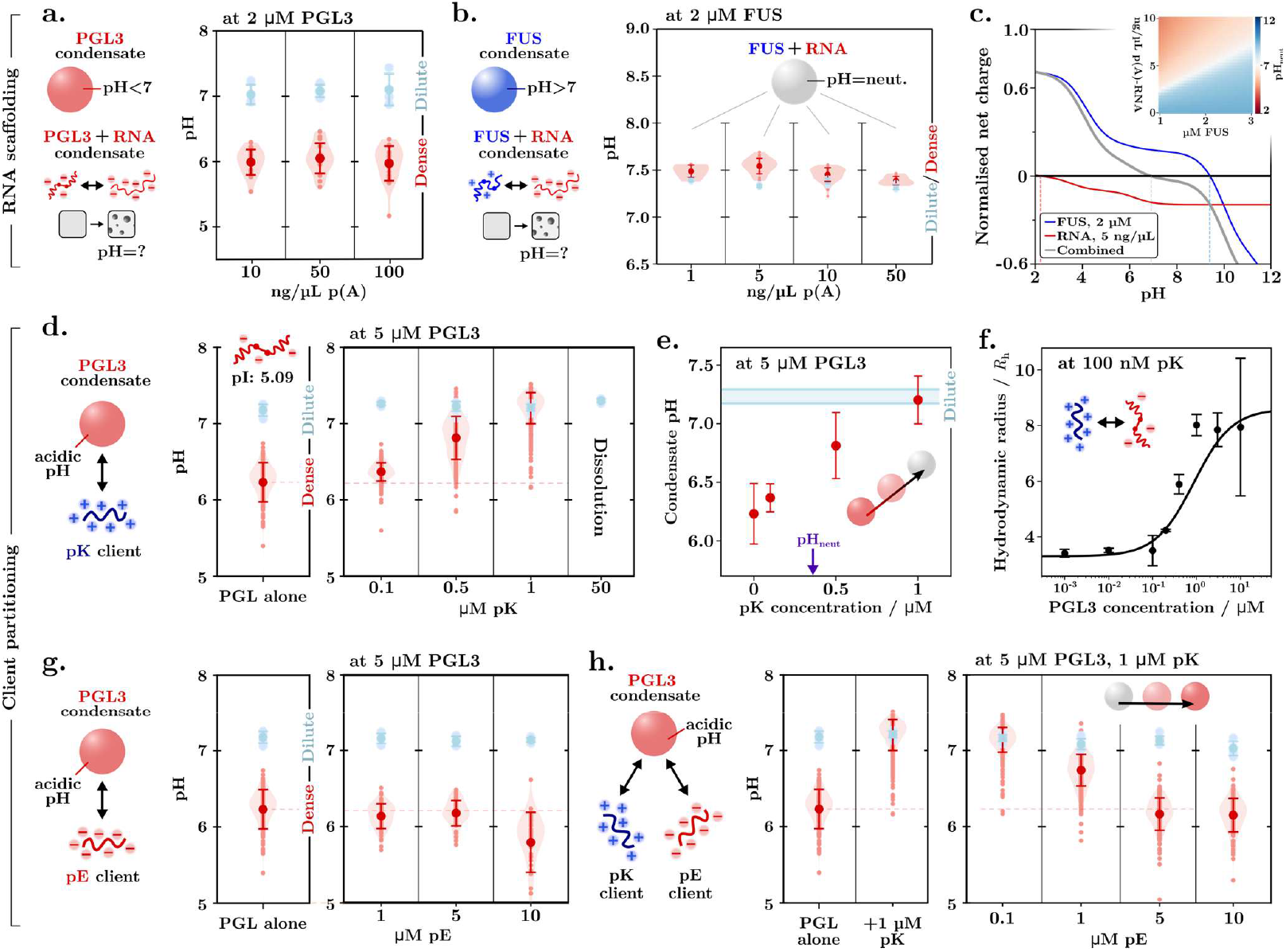
Condensate pH can be dynamically engineered through composition. **(a)** PGL3 dense and dilute phase pH at varying KCl and PEG concentrations. Fixed total protein concentration and buffer condition (2 μM protein, pH=7.3, 50 mM TRIS buffer, 150mM KCl in g and h). Data are presented as mean values ± SD. **(b)** FUS-RNA co-condensate pH at varying p(A)-RNA concentrations. While homotypic FUS condensates display an alkaline dense phase pH, heterotypic FUS-RNA condensates display neutral dense phase pH across a range of conditions. **(c)** Net charge change of FUS with varying pH in comparison to p(A)-RNA and FUS/RNA mixture. The normalisation is performed to account for the concentration difference between RNA and protein followed by bounding to the maximum FUS charge. Inset: heatmap displaying the pH at which neutral net charge is reached for different FUS/RNA compositions. **(d)** Addition of poly-Lysine (pK) to formation of PGL3 condensates and observed dense phase pH values. **(e)** Change in dense phase pH with increasing pK concentration. Blue range: indicates the dilute phase pH. Purple arrow: composition at which the net charge of the pK + PGL3 mixture is predicted to be neutral at physiological pH. **(f)** Quantification of monomer binding between FITC-tagged pK and PGL3. Hydrodynamic radius measurements are performed using microfluidic diffusional sizing. **(g)** Effect of poly-Glutamic acid (pE) to formation of PGL3 condensates and observed dense phase pH values. **(h)** Simultaneous addition of pE and pK to PGL3 condensates. Addition of positively charged pK neutralises the acidic PGL3 condensate pH. Upon increasing addition of pE at constant PGL3 and pK concentrations the condensate pH is shifted back to acidic conditions.

We next set out to study the effect of co-phase separation of the positively charged FUS protein with negatively charged RNA. FUS phase separation under addition of p(A)-RNA abrogates the alkaline pH gradient observed under homotypic conditions (see Fig. 3b). Here, the formation of a pH gradient is no longer required because the heterotypic FUS-RNA interactions enable attraction even at the environmental pH. This decreases the requirement for both highly charged species to form a pH gradient yielding an effective pI of the mixture of macromolecules somewhere between the pI’s of individual species. To quantitively account for this effect, we then consider the charges contributed by both RNA protein as well as accounting for the relative concentration differences in the experiment. Crucially, we find an effective net charge neutralisation of FUS-RNA mixtures in the range of tested RNA concentrations (Fig. 3c). When the RNA concentration is varied, the pH of FUS–RNA condensates remains unchanged, implying stabilization at an electrostatically neutral, fixed stoichiometry. Consequently, altering the overall RNA concentration is not expected to affect the dense-phase composition. Indeed, measurements of the dilute-phase FUS concentration reveal a minimum near the predicted point of electrostatic neutrality, indicating that phase separation is maximized at this optimal FUS– RNA stoichiometry (SI Fig. 12).

### Client-driven condensate pH gradient tuning

Proteins can also passively partition into condensates without triggering phase separation or providing scaffolding. Clients with distinct electrostatic properties, therefore, might enable modulation of the electrostatic repulsion and thus the dense phase pH. To test this, we next subjected PGL3 condensates formed through low ionic strength conditions to addition of highly charged poly-amino acid clients (poly-Lysine, pK and poly-Glutamate, pE). Upon addition of the positively charged pK the acidic pH gradient observed for PGL3 condensates decreased (Fig. 3d), even scaling linearly with pK concentration until effective abrogation of the pH gradient (Fig. 3d, e). Notably, a higher concentration of pK than predicted for charge neutrality was required to achieve neutral condensate pH, likely owing to incomplete pK partitioning (Fig. 3e, purple arrow). Beyond this point, phase separation ceased because stoichiometric binding prevented network formation required for condensation (Fig. 3f).

We then proceeded to also add negatively charged pE to acidic PGL3 condensates. Here, while limited effects were observed, at high pE concentrations the condensate pH decreased even below the conditions formed by PGL3 alone. To study the effects of multiple clients we then also added both pE and pK to PGL3 condensates starting from PGL3 and pK conditions where condensates displayed no pH gradient. Crucially, upon addition of pE the dense phase pH continuously decreased towards the acidic conditions observed for homotypic PGL3 condensates. Thus, our data highlight that client addition can enable precise control over the dense phase pH through altering the effective electrostatic component mixture.

### Multiphasic, multi-pH condensates

We next examined client partitioning in protein-RNA condensates by forming PGL3-RNA condensates, which are acidic in the absence of clients, while adding positively charged pK. This produced multiphase condensates with distinct PGL3- and pK-rich phases (Fig. 4a, b). Binding affinity measurements showed that pK binds p(A)-RNA with much higher affinity than PGL3 (Fig. 4c), explaining the multiphase behaviour: both proteins compete for RNA binding sites, favouring separate pK-rich domains where pK–RNA interactions predominate. Indeed, increasing the pK concentration led to increased numbers of multiphasic structures (Fig. 4d). Using SNARF-4-based pH readouts (Fig. 4d, e), we observed that each phase possessed its own pH level: pK-rich regions were slightly alkaline, whereas PGL3-rich domains were slightly acidic, reflecting their opposing electrostatic properties (Fig. 4f). Thus, condensates not only maintain a pH gradient relative to the dilute phase but can also form multiphasic, multi-pH architectures via segregative demixing (Fig. 4g).

**Figure 4:**
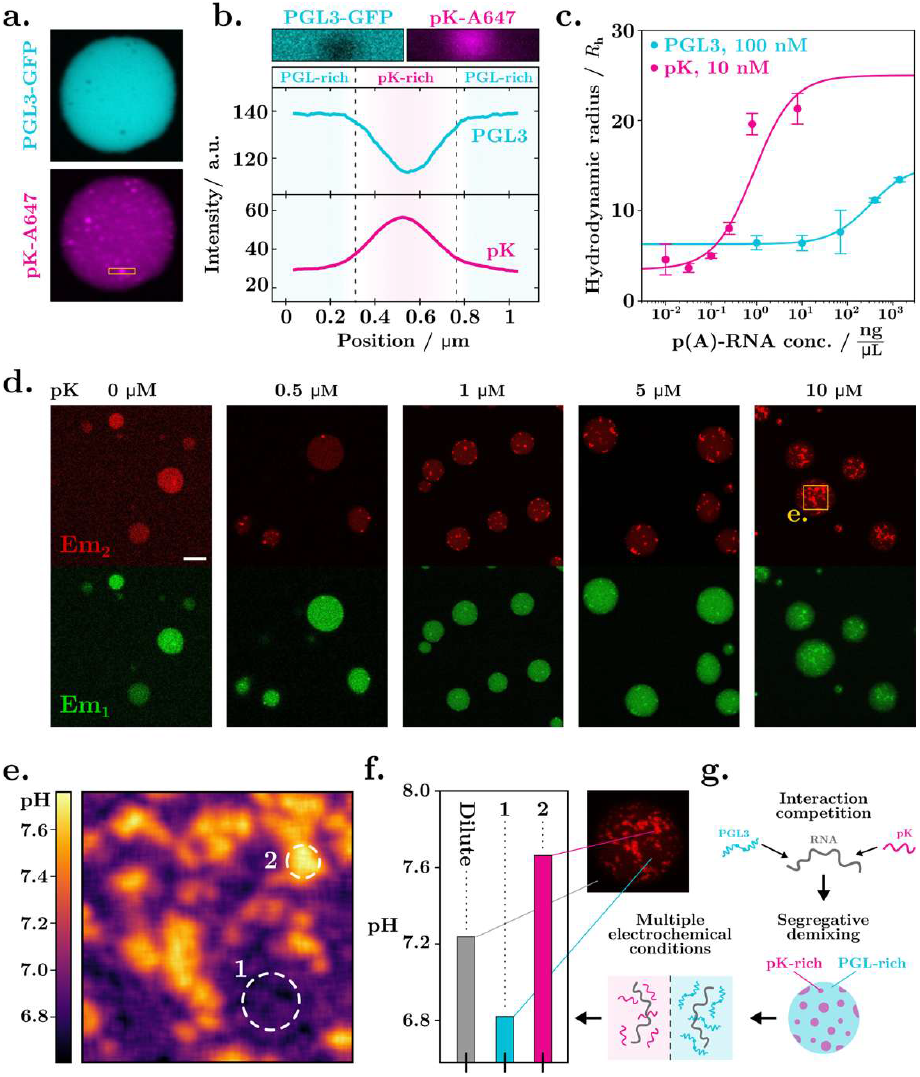
Multiphase, multi-pH condensation of PGL3 with p(A)–RNA and poly-lysine (pK). **(a)** Confocal images of PGL3, pK, and p(A)-RNA condensates, with covalent fluorescent labels on PGL3 (cyan) and p(K) (magenta). **(b)** Fluorescence intensity profiles across a multiphasic condensate sub-feature, indicating distinct pK- and PGL3-rich domains. **(c)** RNA binding curves for PGL3 and FITC-tagged pK, obtained by monitoring hydrodynamic radius changes via microfluidic diffusional sizing. **(d)** SNARF-4F–based pH imaging of PGL3–pK–RNA condensates at different pK concentrations (scale bar: 5 µm). **(e)** Spatially resolved pH map of a multiphasic condensate from panel (d), highlighting alkaline (pK-rich) versus acidic (PGL3-rich) regions. **(f)** Quantitative comparison of local pH values in distinct multiphase sub-features relative to the dilute phase. **(g)** Schematic illustrating competitive RNA binding by PGL3 and pK, which drives the formation of segregated condensate domains with unique electrochemical microenvironments at equilibrium.

### Phase separation-prone proteins in general are highly charged

Our data suggest that proteins with distinct physico-chemical properties can set up diverse pH gradients. Hence, we next sought to characterise the electrochemical diversity of the naturally occurring proteins by plotting the pI distribution of the human proteome (n = 20324). This yields a distinct bimodal shape (see Fig. 5a) highly devoid in proteins with low net charge around physiological pH with only 8.5% of all human proteins displaying an isoelectric point between 7-8. Here changes to the pI based on folding effects were not considered further after the overall proteome pI distribution was observed to only change slightly via application of the the PypKA software^41^ (SI Fig. 13). While this charge bias is likely associated with guaranteeing cytosolic solubility, it means that proteins in general are highly charged. The bimodal distribution is conserved through expression, as shown by mass spectrometry (MS) quantification of proteins present in the U2OS cell lysate^42^ (SI Fig. 14), suggesting that the available intracellular proteins present a similar charge diversity. In the context of phase separation this a potentially widespread requirement for adjusting dense phase repulsion through pH gradients.

**Figure 5:**
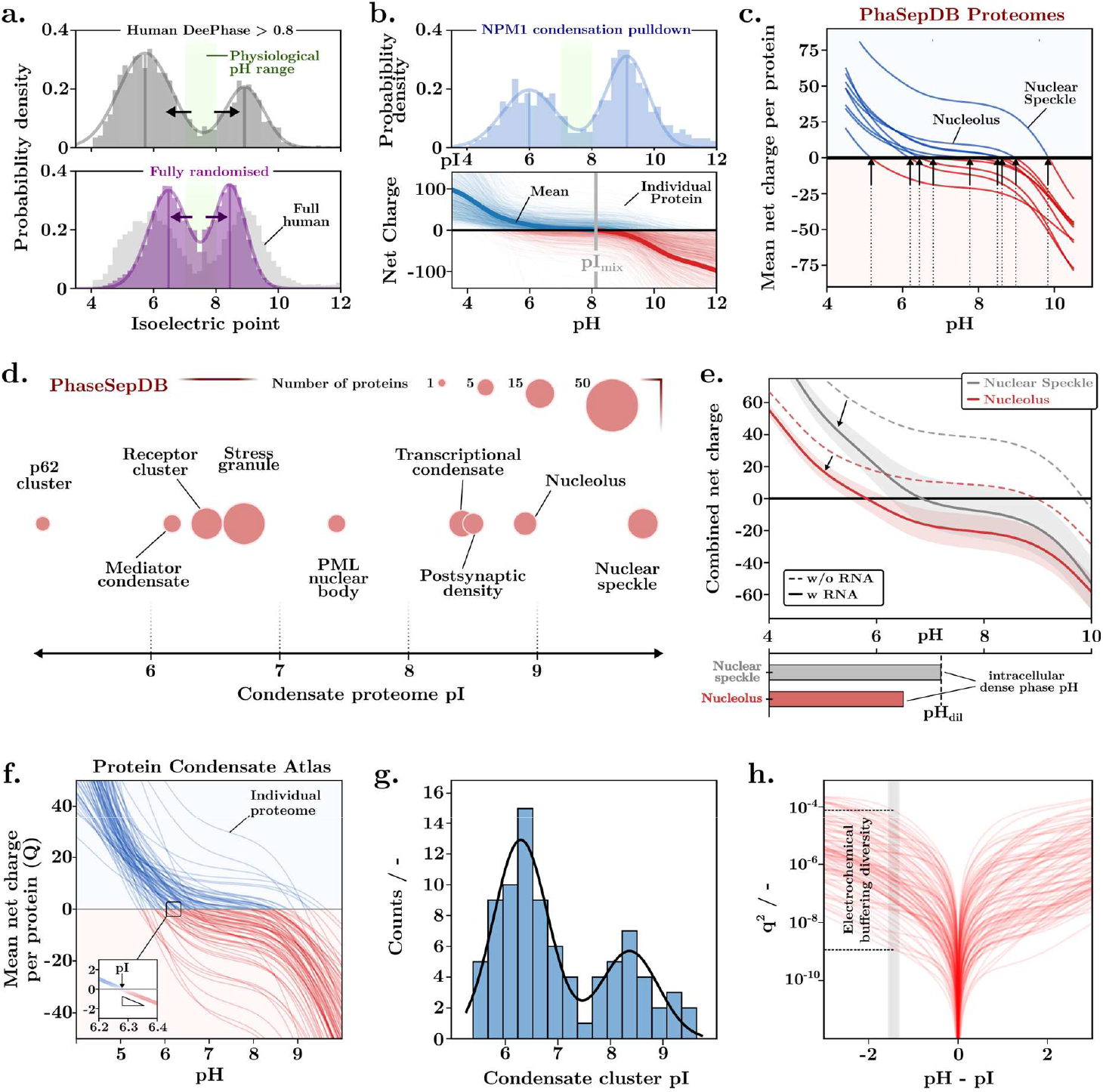
Generic protein charge bias translates to intracellular condensate pH gradients and massive electrochemical diversity of human condensates. **(a)** Isoelectric point distribution of highly phase separation-prone proteins as quantified using DeePhase Scoring (DeePhase Score > 0.8)^43^ (top) compared to fully randomised sequences and the human proteome (bottom). **(b)** Isoelectric point distribution of quantitative proteomics data obtained from NPM1 condensation in cellular lysate from ^44^ (top). Net charge change with pH of the full set of individual proteins obtained from (b) compared to the mean. The point at which the mean net charge curve crosses the x-axis is referred to as the condensate mixture pI (bottom). For correction for the relative abundance of proteins see SI Fig. 18. **(c)** Visualisation of condensate proteome net charge change with pH for complex condensate proteomes obtained from PhaSepDB^45^. **(d)** Comparison of condensate proteome mixture pIs of complex condensate proteomes as obtained via evaluation of protein sequences from PhaSepDB^45^. **(e)** Comparison of nucleolus and nuclear speckle net charge density square profiles and correlation to observed *in vivo* pH gradient from ^26^. Dotted lines show the average net protein charge calculated without RNA. Solid lines incorporate a 7.5% RNA mass fraction, with shaded regions indicating the 5–10% range, consistent with published nucleolar fractionation data^46–48^. **(f)** Schematic representation of the Protein Condensate Atlas from ^49^ used to evaluate the theoretical range of electrochemical properties presented by complex condensates. **(g)** Histogram of condensate cluster pIs obtained for all 85 condensate clusters from the Protein Condensate Atlas. **(h)** Comparison of the net charge density square for the Protein Condensate Atlas. Grey box: highlighting the range of q^2^ values of different condensate cluster at similar pH – pI.

We then sought to understand this charge bias specifically for phase separation-prone proteins. We identified a subset of 3950 proteins with high homotypic phase separation propensity from the human proteome by applying the predictor DeePhase^43^ (Score between 0-1; Threshold > 0.8). This subset of proteins yields a similar pI distribution compared to the full human proteome (see Fig. 5a). Hence, even phase separation-prone proteins are, in general, highly charged (7 < pI < 8: 7.2 % for PS-prone proteins, 8.5 % full human proteome) meaning a large variety of pH gradients could be accessible through phase separation. A disparity in the probability density between acidic and basic pI values can potentially be attributed to a bias of DeePhase for homotypic phase separating proteins as high pI proteins are more likely to engage in protein-RNA interactions. Given the potential importance of the observed charge bias to condensate biology, we set out to understand the origins of the bimodal protein pI distribution further. The distinct shape appears to trace back to the physico-chemical properties of the amino acids themselves, as none of the naturally charged amino acids reach charge neutrality at physiological conditions (see SI Fig. 15), and neither will most combinations of these building blocks. Accordingly, independent species such as yeast (*Saccharomyces cerevisiae*) or even extremophiles (*Acidithiobacillus thiooxidans, Alkalihalobacillus clausii*), that face a necessity to adjust to much harsher environmental conditions, display a lack of proteins with 7 < pI < 8 (SI Fig. 16). Even when considering randomised sequences the bimodal pI distribution is conserved (see Fig. 4a, SI Fig. 17). The probability distribution of human proteome amino acid abundances, however, further creates a bias towards increasing sequence net charge (SI Fig. 17). In summary, because proteins inherently are biased towards high net charge due to the properties of the natural amino acids, neutralizing charged repulsion during condensation likely emerges as a generic feature.

### In silico evaluation of the electrochemical properties of complex condensates

In more complex environments, condensates commonly recruit numerous client proteins, all of which could contribute to the electrochemical properties of the condensate. As an example, we first analysed published proteomics data of a NPM1 condensation pulldown in cellular lysate, mimicking the composition of granular compartments inside the nucleus^44^. To infer information on the electrochemical properties we first considered the pI distribution of the individual proteins in the condensate proteome (i.e. the collection of sequences recruited into the condensate). While the scaffold in NMP1 has an acidic pI, the condensate proteome displays a bias towards proteins with high pI (Fig. 5b, SI Fig. 18). Hence, the electrochemical properties of complex condensates appear to be highly dependent on the set of recruited clients, in agreement with our experimental observations. In fact, the positive charge bias slightly increases when considering the relative abundance of proteins in the dense phase as the fraction of proteins with pI > 7 shifts from 63% to 68%. It is unsurprising that client partitioning specificity is heavily biased by the electrochemical properties of the proteins in the dense phase. The formation of a chemical environment favouring, for example, the interaction of positively charged proteins with nucleic acids, is likely to drive further recruitment of sequences with similar physico-chemical properties.

We then proceeded to evaluate the pH response of the condensate proteome by averaging the net charge change with pH of the individual sequences (Fig. 5b, bottom panel). We find an effective condensate mixture pI (pH at which charge neutrality is reached for the proteome) of around 8.5, with relative abundance (pI = 8.7) and non-corrected data (pI = 8.2) closely correlating (see SI Fig 18). This suggests that the approximation of assessing only the properties of the unique sequences recruited functions well as a proxy in this case. We then applied this approach to common condensate systems acquired from PhaSepDB^45^ including nucleoli and nuclear speckles (see Fig. 5c, d). Here, the obtained condensate proteome pI for the nucleolus at 8.9 correlates well with the values obtained for the simplified NPM1 lysate pulldown mimic. Hence, suggesting consistency in the electrochemical property profile for these condensates independently of the underlying methodological identification approach.

We subsequently wanted to understand how the proposed *in silico* electrochmemical reconstitution translates to potential condensate pH gradients *in cellulo*. We turned to published quantification of the dense phase pH of both nucleoli and nuclear speckles as determined via expression of the ratiometric protein-based pH sensing dye pHluorin2^26^. Both nucleoli and nuclear speckle condensate proteomes are characterised by positively charged mixture pIs (Fig. 5d, e), yet significant recruitment of negatively charged nucleic acids is expected. To account for the RNA contribution, we assumed an approximate mass fraction of 5-10% in these complex condensates, consistent with published nucleolar fractionation data^46–48^. When considering both the protein and nucleic acid contributions in this way, we find that the Nuclear speckle proteome is expected to reach charge neutrality at pH’s around 7. The Nucleolus in comparison is on average constituted of fewer, less positively charged proteins, yielding a charge profile which only becomes neutral at pH < 7 when considering both contributions from proteins and RNA. Accordingly, in cells nucleoli create slightly acidic microenvironments while nuclear speckles were shown not to sustain a pH gradient (Fig. 5e). As such, *in silico* evaluation of condensate proteomes can also inform us of the observed electrochemical condensate properties *in vivo*.

### Dynamic electrochemical microenvironments through complex condensation

We next sought to characterise the potential effector range and functional diversity of electrochemical gradients of condensates. Crucially, even the mapped set of condensate proteomes from PhaSepDB display a large diversity of electrochemical properties (Fig. 5c, d). Compared to the positively charged proteomes of nuclear compartments, stress granules, for example, display a slightly acidic mixture pI of 6.8. The full set of the top nine membraneless organelle systems from PhaSepDB even span a range of almost five pH units in condensate proteome pI’s (Fig. 5d). This indicates that condensates present a large range of electrochemical properties *in vivo*.

To investigate the electrochemical diversity of condensates further, we used the Protein Condensate Atlas. The Atlas comprises predictions for condensate systems generated through an unsupervised clustering of a protein-protein interaction network^49^. The clusters are filtered for a high proportion of predicted phase-separating proteins giving a set of 85 condensate cluster proteomes (SI Fig. 19). We subjected these proteomes to quantification of the pH-dependent net charge changes (Fig. 5f), to find mixture pI’s fully covering the range of pH 5 - 10 (Fig. 5g). This suggests that condensates could potentially allow for the formation of a large diversity of dynamic pH microenvironments suited to the specific function of the individual system. The mixture pI distribution closely resembles the double gaussian shape of the human proteome, likely because of protein abundance bias. A disparity in peak heights between acidic and basic pH values could be attributed to favouring predictions of condensates that are less dependent on protein-RNA interactions.

We then also evaluated the pH-responsive q^2^ behaviour for all 85 condensate clusters (see Fig. 5h). Upon centring these profiles at the mixture pI, we find that the q^2^ terms can vary up to five orders of magnitude at ~ 1.5 units below the respective mixture pI. This suggests that condensates will not only be able to set up a range of pH gradients but also display distinct response behaviours to environmental perturbations. The protein net charge change with pH at the mixture pI: –(ΔC/ΔpH)_pI_, for example, displays a variation of a factor of five for systems with pI ~ 6 (SI Fig. 19). Hence, some systems might be able to sustain the charged state of their constituting proteins even upon changes in the environmental pH; other condensate clusters might change the protonation state of the constituting proteins in a switch like manner.

### Condensates as spatially compartmentalised buffers

Our data suggest that condensates maintain pH gradients to minimize electrostatic repulsion and that electrochemical buffer systems arising from condensation may be widespread in nature. To further elucidate the origins of condensation-driven electrochemical buffering, we developed a minimal theoretical model in which a polymer can adopt positive, negative, or neutral charge states within an aqueous, buffered environment (Fig. 6a). This model incorporates a generic attractive interaction that promotes phase separation, alongside electrostatic repulsion among charged states (see Appendix ‘Theoretical Model’ for details). By analysing the Hessian, we identify regions of phase instability (the spinodal). As expected, we found a minimum in the polymer’s critical concentration near a predefined isoelectric point, confirming that phase separation is most favourable when net charge is minimised.

**Figure 6:**
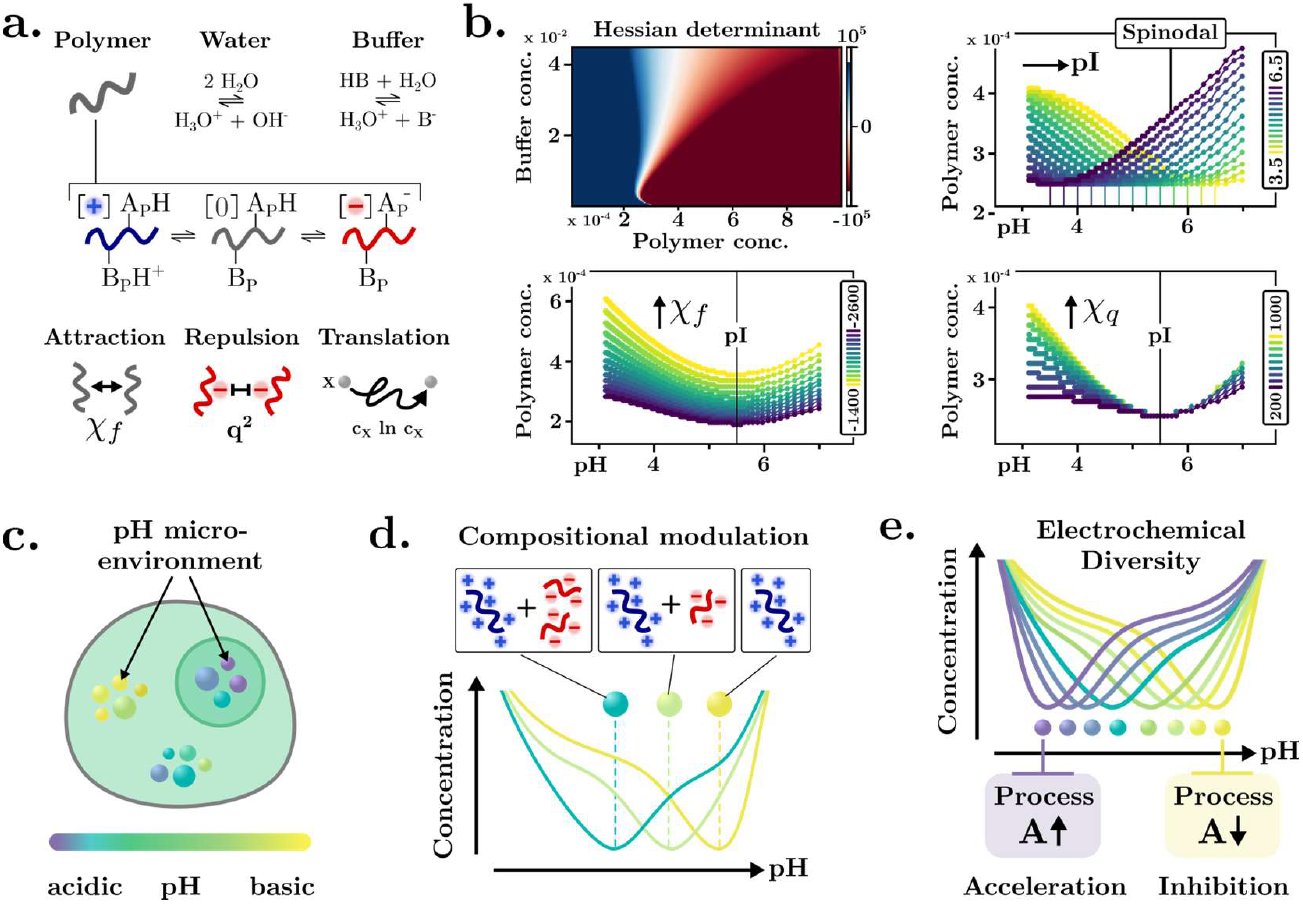
Biomolecular phase separation as a mechanism to generate distinct pH microenvironments. **(a)** Theoretical model set-up to calculate the pH dependent phase behaviour of a generic polymer. **(b)** Prediction of composition-dependent spinodal positions from solving for the theoretical model and variation of key parameters in the pI, attractive Flory and repulsive electrostatic parameter. All concentrations are given in M. **(c)** To minimise electrostatic repulsion biomolecular condensates, sustain distinct pH microenvironments at equilibrium without necessitating membrane enclosure or the presence of active processes. **(d)** Condensates pH and pH-responsiveness can be dynamically fine-tuned through biomolecular properties, compositional changes or complex condensate architectures affecting the overall charge distribution. **(d)** The diversity of electrochemical properties of complex condensates suggests potential for segregation and selective acceleration or inhibition of a range of functional processes, as highlighted visually by the modulation of a generic process A.

Increasing the generic attraction (decreasing χ_f_) shifts the spinodal downward, allowing phase separation at pH values farther from the polymer’s pI for a fixed polymer concentration (Fig. 6b). Conversely, stronger electrostatic repulsion narrows this minimum, requiring higher polymer concentrations to enable separation at pH values offset from the pI (Fig. 6b). In other words, the homogeneous phase becomes unstable primarily near conditions that reduce net electrostatic repulsion, driving the system toward a stable, high-polymer-density regime. In the resulting condensed phase, the pH will then be determined by the underlying acid-base equilibrium to which the high concentration of charged polymer side chains will contribute, while buffer species and other components can also partition differentially. Consequently, condensates effectively function as spatially compartmentalized buffers, forming and sustaining pH gradients at equilibrium.

### Discussion and Conclusion

Effects of pH on the rate of biological and chemical processes, by e.g. regulating enzyme activity or direct catalytic function of H^+^ and OH^-^, are well documented. Membrane bound organelles, for example, commonly display distinct pH conditions aimed at optimising processes relevant to their function, but require energy input to maintain^7–12^. In this study, we reveal that condensation presents a generic mechanism for the formation of pH gradients at equilibrium without necessitating active processes through spatially compartmentalised buffering.

Emergent properties arising from collective interactions have been a point of contention since the initial discovery of protein condensation. Our work details that condensates sustain pH gradients to mitigate electrostatic repulsion between charged proteins in the dense phase. This pH regulation is only possible through the formation of a distinct phase, which creates differential solute partitioning and high local protein concentrations to set-up a distinctly buffered environment from the dilute phase. In protein folding or aggregation, for example, charge neutralisation is similarly required for assembly. However, the electrostatic repulsion commonly has to be overcome by shifts in the pK_a_s of residues in the protein core or the assembly interface^50–55^.

While biomolecular condensation yields drastic increases in viscosity, the molecular scale dynamics remain rapid enough to support efficient reactions^56^. As such, condensates have been shown to affect reaction rates^57–60^ and recent evidence highlights that such condensate pH gradients can modulate enzyme-catalytic outcomes^61,62^ or modulate redox reactions^25^. Besides, such differential ion partitioning within condensates can give rise to transmembrane-like interfacial potentials^63^. The observed large diversity of electrochemical properties presented by intracellular condensate networks (Fig. 6b), therefore, highlights the potential of biomolecular condensation to segregate and optimise functional processes through changes in the emergent electrochemical environment (Fig. 6c). This promise is further expanded by observations that coupling to non-equilibrium phenomena can even drive complex emergent behaviours such as induced microscopic flows^64,65^ pointing to a potential broader functional repertoire. Similarly, exploring how aging or gelation processes modulate these pH gradients and associated functions will present an exciting area for future research.

Our data also highlights that control of the pH gradient arises from the physico-chemical properties of constituting biomacromolecules allowing for dynamic fine-tuning through composition. Hence, controlling nucleic acid or client recruitment are both expected to function as levers of condensate pH gradients. The various dynamic tools available to biological systems to modify the properties of biomolecules, such as post translational modifications^66–68^, such as phosphorylation, could further enable modification of condensate pH gradients. Together such dynamic modulations or multiple coexisting dense phases with distinct internal pH values could provide additional layers of regulation for sequential reactions or biochemical process cascades more generally.

This also has potential consequences to hypotheses that membraneless compartmentalization might have been an important factor in the origin of life^23,69^. Here, shifts in the pH of protocells formed through condensation of protobiomolecules could be a contributor to generating biological functionality and sustaining required pH conditions. Similarly, pH gradients arising from macromolecular density transitions could be leveraged in technological applications, for example by building on the rich history of polymer or coacervate phase transitions, to generate designer condensation systems with optimal electrochemical properties for desired processes.

## Acknowledgements

The authors thank Prof. Rohit Pappu for valuable discussions and feedback. We kindly acknowledge funding from the European Research Council under the European Union’s Horizon 2020 research and innovation program through the ERC grant DiProPhys (agreement ID 101001615) (H.A., C.M.F., D.Q., T.P.J.K.); Global Research Technologies, Novo Nordisk A/S (H.A., T.P.J.K.), Transition Bio (R.S.), the Frances and Augustus Newman Foundation (T.S., E.d.C.) and the Centre for Misfolding Disease.

## Author contributions

H.A., R.S. and T.P.J.K. designed and conceptualised the study. H.A., R.S., T.S., C.M.F., I.B., E.d.C., A.Z.B, and M.O. performed investigations. T.S., K.L.S., T.M.F., S.W., J.M.I.-A., G.K., G.I., S.A., A.A.H., N.L.Z. and T.P.J.K. provided materials and methods. H.A., R.S. and D.Q. analysed the data. H.A. and T.P.J.K. wrote the original draft of the paper. All authors discussed the results, reviewed, edited and contributed to the final manuscript.

## Correspondence

Correspondence should be addressed to Tuomas P. J. Knowles.

## Data availability

The raw data and analysis code underlying this study will be made available upon request.

