## Supplementary Information for "Biomolecular condensates sustain pH gradients at equilibrium through charge neutralisation"

### Materials and Methods

#### Reagents

All reagents and chemicals were purchased with the highest purity available. p(A)-RNA with a molecular weight range from 700 to 3500 kDa was purchased from Sigma Aldrich as lyophilized powder and dissolved into a stock of 10 mg/mL in water before use. 50% w/v PEG (Polyethylene glycol) 8000 Da stock solution was prepared by dissolving PEG pellets (Sigma-Aldrich) in 25 mM TRIS-HCl pH 7.3 buffer solution. The stock solution was stored at room temperature. Histidine and Succinic acid were purchased from Sigma Aldrich. Cy5-NHS ester was purchased from Thermo Fischer and dissolved in DMSO to give a 10 mM stock concentration and stored at -20 °C. SNARF-4F was obtained from Thermo Fisher and subsequently dissolved in DMSO to give a 10 mM stock concentration, which was stored at -20 °C. Insulin was purchased from Sigma Aldrich and dissolved in high pH H/S buffer (50 mM, pH = 9.0) to prevent precipitation at e.g. 5 mg/mL. Insulin glargine was ordered from Sigma Aldrich and dissolved in water (pH = 4, HCl adjusted) with 2 (w/v)% glycerol, 0.2 mM ZnCl and 25 mM m-cresol. Poly-Lysine (pK, 30-70 kDa), Poly-Lysine-FITC (30-70 kDa) and Poly-Glutamic acid (pE, 15-50 kDa) were ordered from Sigma Aldrich and subsequently dissolved in the respective buffer used for the specified experiment.

#### PGL-3 protein expression

PGL-3-6xHis-mEGFP purification was performed according to a published protocol[1]. Untagged PGL-3 was then obtained by cleaving off 6xHis and mEGFP tags from the C terminus of PGL-3-mEGFP-6xHis protein via usage of 6xHis-tagged TEV protease. Subsequent incubation with Ni-NTA Agarose resin allowed for removal of 6xHis-tagged mEGFP and TEV protease. Further purification of untagged PGL-3 was performed via size-exclusion chromatography using HiLoad 16/60 Superdex 200 column (GE Healthcare) in 50 mM HEPES pH 7.4, 0.3 M KCl, 1 mM DTT. The protein was stored at -80°C followed by aliquoting and flash freezing in liquid nitrogen.

#### FUS protein expression

FUS wildtype, EGFP and SNAP tagged variants were expressed as reported previously in an insect cell expression system[2,3]. After purification, the proteins were stored in 50 mM TRIS buffer (pH 7.3), 750 mM KCl, 1mM DTT, 5% glycerol and at a total protein concentration of 20 µM and 70 µM for wildtype and SNAP variants, respectively.

#### Histidine and succinic acid buffer system preparation

To give the pH = 3.5-9 buffer system (H/S buffer), histidine and succinic acid were first both dissolved together in dd$H_2O$ at 200 mM each. This H/S buffer stock was then split in two equal parts which were then individually adjusted to pH 9 and 3.5 using concentrated HCl and NaOH, respectively. The obtained low pH and high pH H/S buffer stocks were then mixed at different ratios to give volume fractions of pH = 9 H/S buffer from 0-1 in 0.1 steps. These samples were then subjected to measurements using pH probes to give a calibration curve (SI Fig. 1). This calibration curve was subsequently applied in future measurements to directly calculate the pH from the volume fraction of the pH = 9 H/S buffer stock. Furthermore, from the calibration curve stocks at desired pH values could be prepared by mixing pH = 9 and pH = 3.5 H/S buffer stocks at the corresponding volume fractions. All H/S buffer stocks were stored at 4 °C.

#### Fabrication of microfluidic devices

Microfluidic devices were designed using AutoCAD software (Autodesk) followed by printing on acetate transparency masks (Micro Lithography Services). The replica master was obtained via standard soft-lithography steps using spin-coating of SU-8 photoresists (MicroChem) onto polished silicon wafers.[4] Typically, SU-8 3050 was applied to achieve a device height of approximately 100 µm. After UV exposure, utilising a custom built LED-based apparatus[5], the precise heights of the features were measured using a profilometer (Dektak, Bruker). Devices were then produced in polydimethylsiloxane (PDMS). PDMS (Dow Corning) was mixed in a 10:1 (w/w) ratio with curing agent (Sylgard 184, Dow Corning) and poured onto the master, followed by degassing and baking for 1.5 h at 65°C. The PDMS was then removed from the master and punched using biopsy punches to generate inlet holes, after which the slab was bonded onto thin glass slides using oxygen plasma surface activation (Diener electronic, 40 % power for 30 s).

#### Continuous pH phase boundary mapping

For the microfluidic experiment four aqueous solutions containing peptide, dd$H_2O$ and both high and low pH H/S buffer stocks (50 mM, pH = 3.5 and 9, respectively) as well as an oil solution for droplet generation (HFE-7500 mechanical oil containing 1.2% Bio-RAN) were used. Here, the peptide was tagged with Cy5 NHS-ester dye (1:100 molar ratio) using a previously established procedure[6] before application of to the microfluidic chip the stock. Both H/S stock solutions were then individually supplemented with either Alexa Fluor 488 or 647 free dye at a concentration of 5 µM, to enable tracing of the high and low pH buffer fraction in each droplet. The solutions were then loaded into four separate inlets on the microfluidic chip using pressure control pumps (LineUp Flow EZ, Fluigent). The aqueous solutions were first contacted before encountering the droplet junction, where droplets were formed by a constant oil flow of 100µl/h. To scan chemical phase space the total flow rate of both H/S buffer stocks combined were kept constant while varying their flows individually between 5 and 50µl/h and ensuring a final experiment buffer concentration of 50 mM typically. Similarly, the combined flow rate of peptide and $ddH_2O$ flow rate was kept constant while varying both flows individually between 5 and 50µl/h at much shorter intervals compared to the H/S buffer flow rate variations. Droplets were subsequently incubated for 3 minutes on chip followed by imaging under continuous flow using an openFrame epifluorescent microscope (Cairn Research) with a 10x 76 air objective (Nikon CFI Plan Fluor) and equipped with a dichroic filter set (Cairn Research) allowing simultaneous imaging of three wavelengths (488nm, 546nm, 647nm). Crosstalk was accounted for by imaging the stock solutions individually. These images were subsequently analysed using a custom-written Python (Python version 3.9.7) script, enabling both droplet detection and phase separation classification[7]. Phase separation classification was assigned as mixed = 0, phase separated = 1. The phase separation probability $P_{PS}$ was then calculated by averaging the 0/1 phase separation classification of neighbouring data points (radius of 3% of the maximum axis limits) for each individual data point or by application of binning based on established literature algorithms8,9 (see SI Fig. 2). To calibrate, microdrops filled with only one stock solution at a time are generated and subsequently images in the corresponding fluorescent channel. This allows for correcting the illumination profile by dividing by the average intensity of many frames and then fit a gaussian to the intensity histogram of the calibration images to obtain an intensity value which corresponds to the stock solution's concentration (SI Fig. 3). The same process is applied for a background image (droplets with no corresponding dye), and a calibration curve of droplet intensity to concentration is generated. The process is repeated for all fluorescent solutions. For mapping intensity to the pH of the droplets, normalised intensities in the respective high and low pH fluorescent wavelengths were used to establish the volume fractions of high and low pH buffers.

**Sample preparation and phase separation conditions**

Phase separation was exclusively induced in vitro by gently mixing components in 500µL Eppendorf tubes, where the specific conditions investigated are reported for each specific dataset. To probe phase separation of FUS under varying concentrations of TRIS, $ddH_2O$ was first added to the Eppendorf tube followed by addition of KCl (1M in $ddH_2O$) to give a constant final concentration of 150 mM considering and addition of TRIS (1M, pH = 7.3) to obtain a range of 20-200 mM final concentration. Lastly the protein (70 µM, 750 mM KCl, 50 mM TRIS buffer pH = 7.3) was added to yield a final concentration of 1 µM at a total volume of 10 µL. Fluorescence was then observed via an inverted microscope (OpenFrame, CairnResearch, Faversham, UK) set up with a 20x air objective (Nikon, Surbitton, UK) combined with appropriate filters (Laser2000, Huntingdon, UK). Imaging was then performed with a high sensitivity camera (Prime BSI Express sCMOS, Photometrics, Tucson, AZ, USA).

Insulin dilute phase concentration measurements under varying buffer concentration conditions were performed by first diluting high and low pH H/S buffer (200 mM, pH = 9 and pH = 3.5, respectively) individually to the desired concentration such as 1/10 in ddH2O for a final concentration of 20 mM. Insulin stocks were then prepared at 5 mg/mL in high pH H/S buffer at the same buffer concentration. Samples were then prepared by mixing the Insulin stock with high and low pH H/S buffer at 120 µL, 105 µL and 75 µL each to give a final ratio of 1/3 of low to high pH H/S buffer with a pH of 6.4 (see SI Fig. 1 for H/S buffer pH calibration) and 300 µL total volume at 2 mg/mL final insulin concentration. Samples at varying buffer concentrations were then simply generated in the same way by mixing insulin and H/S buffer stocks with the desired buffer concentration. Experiments at a final insulin concentration of 1 mg/mL were prepared in the same way using 2.5 mg/mL insulin stocks. Insulin dilute phase concentration measurements under varying pH conditions were performed by mixing Insulin stock (10 mg/mL, 100 mM H/S buffer, pH = 9) at a 1/5 ratio with high and low pH H/S buffer stocks (100 mM, pH = 9 and pH = 3.5, respectively) combined to yield the desired pH. The samples were then subsequently centrifuged at 10000 RCF for 10 min followed by decanting of the supernatant and measurement of the intrinsic supernatant absorbance at 280 nm.

##### SNARF-4F pH calibration and readouts

SNARF-4F calibration samples were prepared by mixing high and low pH H/S buffer (100 mM) at varying volume fractions to give stocks in the pH range between 3.5-9, followed by addition of SNARF-4F (in DMSO, 10 mM) to a final concentration of 20 µM. These samples were then first used to characterise the changes in the SNARF-4F emission spectra using Cary Eclipse Fluorescence Spectrophotometer (Varian, Inc.) with an excitation wavelength of 488 nm and scanning emission between 520nm and 730nm with a rate of 120nm/minute and collecting intensity every nm. The excitation slit used was 5nm and emission slit 10nm. Then, to establish the confocal imaging approach 10 µL of the SNARF-4F calibration samples were deposited on clean 24 × 50 mm No. 1 cover glass slides (VWR) and imaged via Leica Stellaris 5 confocal microscope equipped with 63X oil immersion objective (Leica HC PL APO 63×/1.40 Oil CS2, NA 1.4). Here, high-resolution images were taken (2048 x 2048 pixels) by excitation at 488 nm and acquiring at 580±10 nm and 650±10 nm simultaneously. The laser power was kept at 2% and the detector gain was set to 60%. The images obtained at extreme pH conditions (pH = 3.5 and 9) were then simply used to set up the pH reference calibration according to the supplier manual, where $Em_1 = \lambda_1 = 580\pm10$ nm and $Em_2 = \lambda_2 = 650\pm10$ nm. Phase separated samples for dense and dilute phase pH measurements using SNARF-4F were acquired in the same way and prepared by mixing components in 500µL Eppendorf tubes. The specific conditions investigated are reported for each specific dataset and SNARF-4F was consistently applied at a final concentration of 20 µM. Analysis of the dense and dilute phase pH was then performed by feature segmentation of dense phase compartments into image masks using the *ilastik* segmentation toolkit[10,11] (see SI Fig. 6). The dilute phase $Em_1$ and $Em_2$ intensity was then calculated as the average non-dense phase intensity. The $Em_1$ and $Em_2$ intensity of each individual dense phase compartment were calculated by averaging over all pixels associated to only that specific dense phase compartment.

##### Hydrodynamic radius determinations using Microfluidic Diffusional Sizing

Microfluidic diffusional sizing was performed using the Fluidity One-M instrument (Fluidic Analytics) based on previously published operation principle[12]. Briefly, 4 µL of sample at 100 nM protein concentration (unless specified otherwise) were added to the chip after the auxiliary channels were primed with buffer (pH=7.3, 50 mM TRIS buffer, 150mM KCl). Samples were then measured in triplicates based on detection in the Alexa488 (PGL3-GFP, pK-FITC).

##### Mutliphasic codensate imaging

Multiphasic PGL3, pK and p(A)-RNA condensates were formed by mixing X µM of PGL3-EGFP, X µM of pK tagged with Alexa647 and X ng/uL RNA in an Eppendorf tube at a total volume of 20 µL and buffer conditions of 150 mM KCl and 50 mM TRIS. The pK labelling was performed according to a previously established NHS-Ester covalent conjugation protocol[6]. Samples were then imaged after 5 minutes of incubation by deposition on clean 24 × 50 mm No. 1 cover glass slides (VWR) and imaged via Leica Stellaris 5 confocal microscope equipped with 63X oil immersion objective (Leica HC PL APO 63×/1.40 Oil CS2, NA 1.4), taking high-resolution images (2048 x 2048 pixels).

##### Insulin dilute phase concentration measurements

Insulin phase separating samples preparing 1 mL samples at specified conditions, for example 2 mg/mL Insulin in 50 mM H/S Buffer at pH = 6.4, by addition of Insulin from a dissolved pH = 9.0 H/S buffer stock followed by incubation over 10 minutes. Subsequently, samples were centrifuged for 5 minutes after which the supernatant was decanted and dilute phase concentrations were determined using 280 nm absorbance measurements.

##### FUS dilute phase concentration measurements

Given the low material volumes available, FUS dilute phase concentrations were performed by a previously published flow-cell coupled scanning confocal approach[13,14]. Briefly, FUS samples are first prepared at specified conditions in Eppendorf's at a total volume of 10 µL and following 30 minute incubation microfluidic chips were flushed with the sample by applying negative pressure. Sample time traces were then recorded using a home-built confocal setup equipped with picosecond-pulsed 485-nm laser sources as well as a 60x water-immersion objective (CFI Plan Apochromat WI 60x, NA 1.2, Nikon). The dilute phase concentration was then extracted from the time trace as the baseline signal given the large excess of the dilute phase volume fraction compared to the dense phase.

##### Sequence-based physico-chemical property calculations

Physico-chemical properties of individual protein sequences were calculated directly from the sequence using tabled properties of the individual amino acids by applying the *Bio.SeqUtils.ProtParam* python package functions *.charge_at_pH()* and *.isoelectric_point()*. The electrochemical reconstitution of complex mixtures of sequences was performed by first calculating the charge dependent pH profiles of the individual sequences followed by summation and normalisation for the total number of protein sequences. Here, the mixture isoelectric point was simply determined as the point were this effective mixture charge profile reaches zero net charge. The sequence charge density q was determined by normalising the net charge at a given pH with the number of residues. The sequence hydrophobicity was evaluated using Kyte-Doolittle scaling. Species proteomes were downloaded from uniprot. pH-dependent RNA charge calculation was performed using a simplified, site-based approach where each nucleotide in the RNA chain is treated as a combination of a nucleobase and a phosphate group. Each nucleotide is considered as comprised of a nucleobase and a phosphate group. For nucleobases (A, C, G, U), approximate pKa values are assigned, and the fraction of protonation was determined via the Henderson–Hasselbalch equation, thereby estimating any positive charge contributions. Simultaneously, each phosphate group is modelled with two sequential deprotonation steps—each with its own pKa—resulting in net negative charge as deprotonation increases. Finally, the contributions from all nucleobases and phosphate groups across the RNA sequence are summed to produce a theoretical net charge at a given pH.

**Evaluation of impact of folding on predicted pI using PypKA**

To consider folding induced shifts on the calculated protein pI PypKA was applied[15], which takes as input a 3D PDB structure file, and uses a mixed Poisson-Boltzmann and Monte Carlo approach to attain a isoelectric point accounting for the protein's folded state. Application to FUS and PGL3 (SI Table 2) showed limited differences between sequence based and structure-corrected isoelectric point indicating that in these cases simplified sequence analysis is sufficient to capture the electrostatic properties of these proteins. For folding effect testing at scale a similar comparison was performed for a subset of 575 proteins obtained by filtering for SwissProt/UniProt Human proteins and focusing on proteins where full length PDB chains were available (SI Fig. 13). No statistical significance between the two methods (Wilcoxon Signed-Rank Test: p-value = 0.14) were observed while only 15% of proteins have an absolute error greater than 1.5 pI unites when not accounting for the protein's folded state.

**Generation of randomised proteomes**

To obtain information of the phyisco-chemical property distributions of randomised proteomes, proteins were generated by randomly concatenating individual amino acids at given probability of occurrence for the individual amino acids and total protein length. For the set of fully randomised sequences an even probability of occurrence was assumed for the individual amino acids and proteins were constrained to 560 amino acids in total length, corresponding to the average protein length in the human proteome. To obtain a 'humanised' random proteome the probability of occurrence for the individual amino acids was amended to be the same as that of the human proteome and similarly sequences were generated at lengths corresponding to the human proteome length distribution. In each case a total of 500000 sequences were generated and subsequently evaluated for their physico-chemical properties.

**PhaseSepDB condensate proteomes**

PhaseSepDB v2 data was loaded and processed in Python where only 'Homosapien' entries were used[16]. For each membraneless organelle (MLO) in the database, a record of which proteins had this MLO annotation were recorded. Taking the MLOs with 9 most common annotations, the physico-chemical properties of the MLO proteome were evaluated as outlined in above. MLOs with fewer annotations were deemed potentially unreliable to apply this method.

**Condensate Atlas Usage**

Protein clusters from the Condensate Atlas were used if more than 50% of the involved proteins were deemed a 'scaffold' (high homotypic LLPS prediction score) as outlined in [17]. Then, physico-chemical parameters such as charge, and mixed isoelectric point of the condensates were calculated as outlined above.

**Reduced tie line approach in pH chemical space**

To interpret the dilute phase band gradients obtained via the microdroplet pH scanning approach, we first remark that the construction of these bands depends on mass balance, and in the present case the conservation is to be applied to the difference in total concentrations of H+ ions and OH- ions, regardless of whether they are in the free solution state or bound to the buffer, or each other. Mass conservation works for this difference because dissociation of them from buffer molecules does not affect their total concentrations, and dissociation of water releases equal amounts of both to the solution so the difference does not change. The dilute phase bands then indicate how this differential concentration, i.e. the difference between total $[H^+]$ and total $[OH^-]$, is partitioning across phases. To make the connection between the differential concentration and pH, we note that there exists a highly non-linear but monotonic mapping between the two, so each pH on the x-axis of the pH phase boundary corresponds to a particular value of the partial $H^+/OH^-$ total concentration difference, with higher pH corresponding more $OH^-$ compared to $H^+$. A positive dilute phase band gradient at the low-pH section then means the dense phase total $[OH^-]$ - $[H^+]$ is higher than the dilute phase, translating to a higher pH in the dense phase and vice versa for the high-pH section.

#### Supplementary Tables

**Supplementary Table 1:** Sequence comparison between the peptide heterodimers insulin and insulin glargine. Sequence mutations in purple and additions in blue.

| **Protein** | **Chain** | **Sequence** |
|---|---|---|
| Insulin | A | GIVEQCCTSICSLYQLENYCN |
| | B | FVNQHLCGSHLVEALYLVCGERGFFYTPKT |
| Insulin glargine | A | GIVEQCCTSICSLYQLENYCG |
| | B | FVNQHLCGSHLVEALYLVCGERGFFYTPKTRR |

**Supplementary Table 2:** Changes in the pI when considering folded features with PypKA[15] for PGL3 and FUS.

| Protein | Sequence pI | Folded pI | pI error |
|---|---|---|---|
| PGL3 | 5.09 | 4.71 | 0.38 |
| FUS | 9.40 | 9.32 | 0.08 |

#### Supplementary Figures

**Supplementary Figure 1: Histidine and succinic acid linear volume fraction pH buffering system.** **(a)** Molecular structure of histidine and succinic acid including individual relevant $pK_a$s. **(b)** Mixing scheme for generating the buffering stocks by adjusting the pH of mixed histidine and succinic acid stocks. **(c)** Correlation between the measured pH and the volume fraction of pH = 9 H/S buffer stock. **(c)** Salt dependence of the Histidine / Succinic acid buffer system.

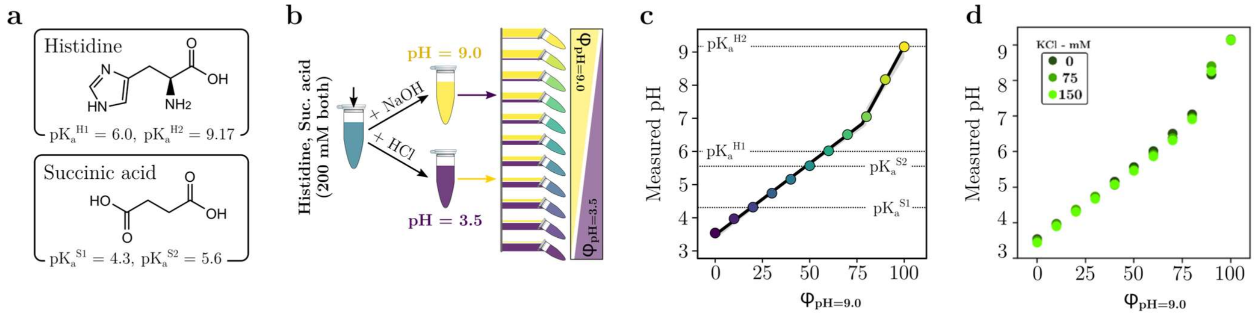


**Supplementary Figure 2: Data processing of microdroplet phase boundary mapping approach.** **(a)** Raw data obtained from droplet based phase boundary mapping. The density of data points (N = 75021) entails that simple scatter plotting is not visually representative of the results obtained given randomized overlapping of individual data points. **(b)** Grid-based averaging then allows to obtain representative visualisation of the data by considering the properties of all data points within a defined bin. Bin sizes were chosen based on established literature algorithms[8,9]. **(c)** Schematic explanation of the visualisation of dilute phase contours. Raw data are first averaged based on neighbouring data points in a defined radius (typically 3% of the axis scale) to generate a representative readout from the large data density. Dilute phase contours are then obtained by repeated sectioning for threshold dilute phase values.

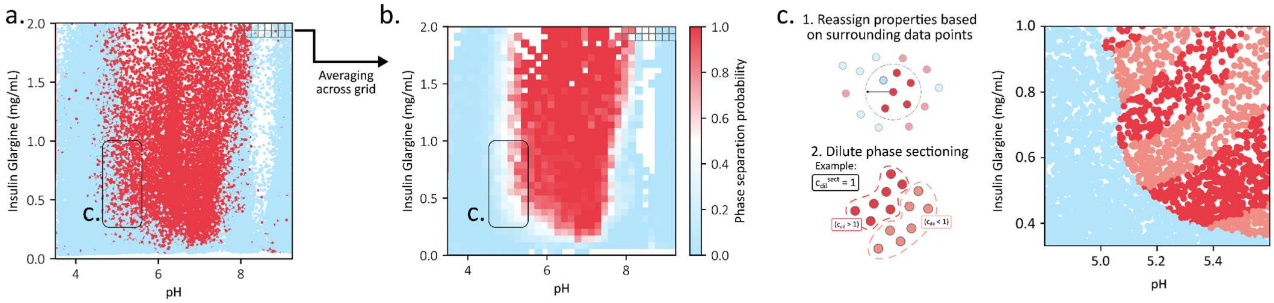


**Supplementary Figure 3: Calibration and correction in droplet microfluidic mapping.** Images of microdroplets containing one of the stock solutions only are first corrected for the Illumination profile (see Raw image vs Ill-pof corrected). Intensity histograms are then fitted to generate concentration calibrations enabling back calculation of the concentration in each individual droplet. This process is subsequently repeated for all fluorescently labelled solute streams.

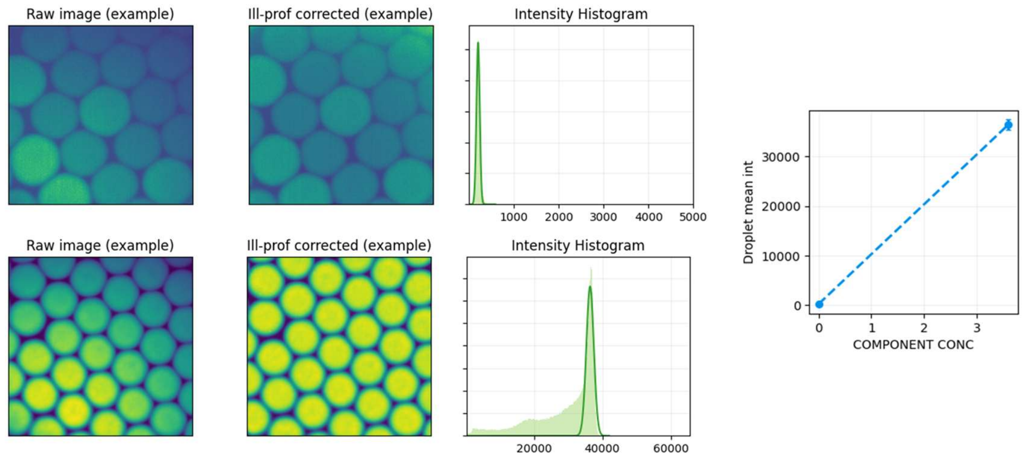

**Supplementary Figure 4: Sequence and phase behaviour comparison of insulin and insulin glargine.** **(a)** Schematic representation of the sequence changes from insulin to insulin glargine. **(b)** pH-responsive phase boundaries for both peptides including indication of the boundary half-width. **(c)** Sequence hydrophobicity profiles for insulin and insulin glargine. **(d)** Sequence net charge profiles for insulin and insulin glargine.

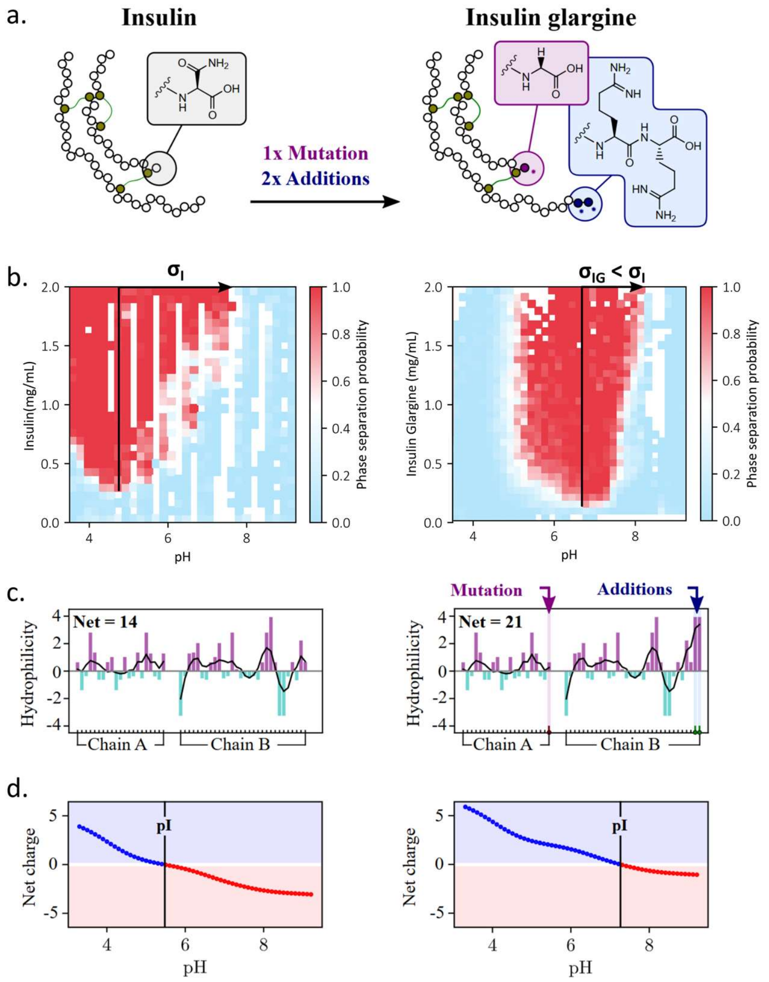


**Supplementary Figure 5: pH-response analysis of insulin phase separation.** **(a)** Change in the insulin dilute phase concentration with varying pH using the histidine and succinic acid buffer system at 2 mg/mL total insulin concentration and constant H/S buffer concentration of 100 mM. The dilute phase concentration is minimized around the isoelectric point indicating that most protein is being converted into the dense phase around the pI. **(b)** Visualisation of the 2D reduced tie line gradients for insulin. Reduced tie line gradients are displayed using red shaded bands as generated by sectioning for the dilute phase concentration obtained in each individual microdroplet. **(c)** SNARF-4F readouts for insulin phase separation at varying pH. Emission maxima readout for both insulin dense (dark blue) and dilute phase (light blue) at varying start pH conditions, highlighting a large variation in the dilute phase but constant dense phase ratio. **(d)** pH dependent q2 profile of unmodified insulin. Approximation of the pH dependent repulsion free energy exerted by insulin in the condensed phase using the sequence charge density (q).

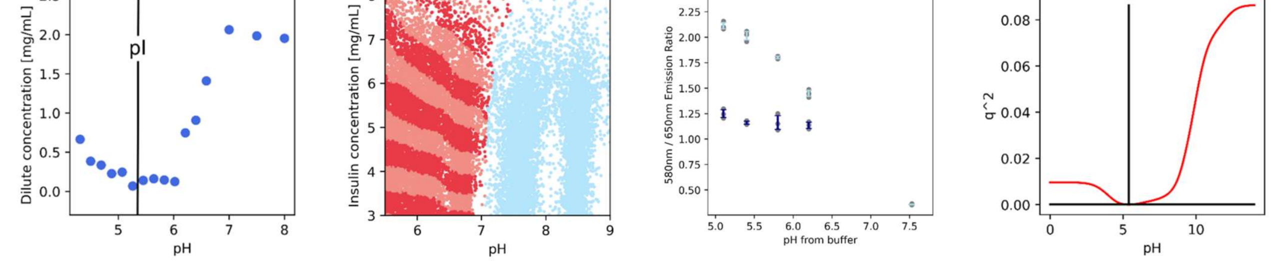

**Supplementary Figure 6: Ratiometric pH-sensing dye SNARF-4FSNARF-4F readout approach (a)** pH dependent emission spectrum shift. SNARF-4F dye emission spectrum post excitation at 488 nm at varying pH, highlighting two emission maxima at around 580 and 650 nm. **(b)** pH readout calibration of SNARF-4F. Change in SNARF-4F intensity ratio of the 580 nm and 650 nm emission maximum as obtained by confocal imaging of SNARF-4F at 20 uM in H/S buffer at varying pH. **(c)** Spatially resolved SNARF-4F based pH readout. Intensity at the 580 nm ($Em_1$) and 650 nm ($EM_2$) emission maxima, emission maxima ratio and pH conversion of SNARF-4F readouts in a spatially resolved manner between dense and dilute phase for insulin glargine at a starting pH of ~ 6 using H/S buffer. **(d)** Exemplary condensate detection mask from SNARF-4f readouts. Confocal image of PGL3 condensates under low ionic strength conditions under the addition of SNARF-4F to read out pH. Image obtained by emission detection in 580 nm (left) and 650 nm (middle) as well as feature segmentation of the individual condensates including annotation (right).

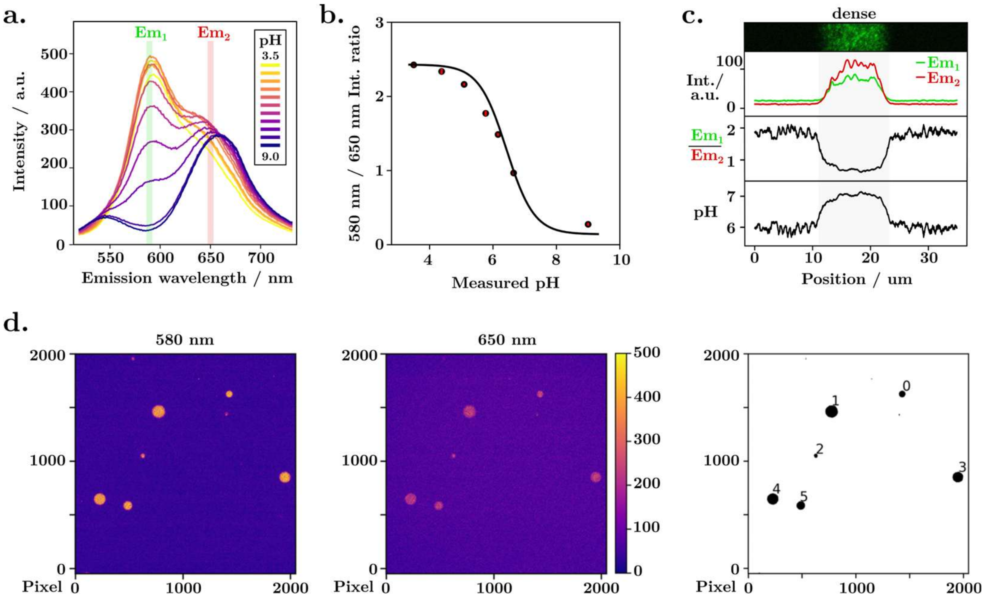


**Supplementary Figure 7: Effect of buffer strength on insulin phase separation.** Insulin dilute phase concentration change with varying H/S buffer concentration at fixed pH = 6.4 at a total insulin concentration of 2 and 1.5 mg/mL.

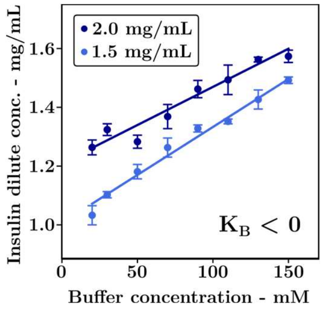


**Supplementary Figure 8: Dominance analysis execution. (a)** pH-responsive phase boundary change with addition of KCL highlighting a data slice used for dominance analysis. **(b)** Change in $c_{dil}$ of insulin glargine with changing total concentration at a fixed pH and salt concentration (see marking box in d.). R - dilute phase response gradient.

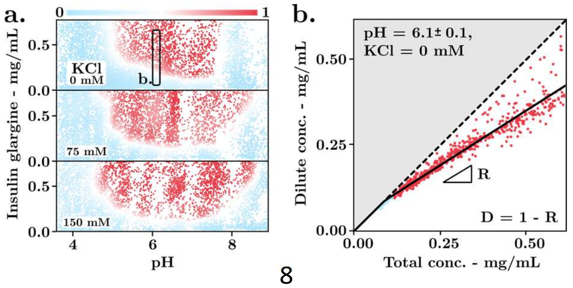

**Supplementary Figure 9: Impact of condensate size on the apparent pH readout.** (Left) SNARF-4F 580 nm/650 nm emission ratio plotted against the number of pixels for each individually analysed PGL3 condensate. Light-shaded background points represent individual data, while magenta symbols and error bars show mean ± SD within each bin of pixel ranges (delineated by dashed vertical lines). (Right) Condensate pH, derived from the average fluorescence ratio, plotted against the effective droplet diameter (assuming spherical geometry).

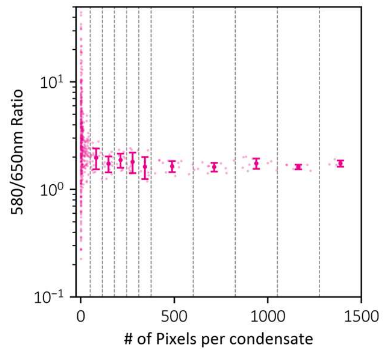


**Supplementary Figure 10: FUS condensate pH under varying conditions. (a)** Schematic of the experimental workflow to probe the effect of various environmental conditions on the obtained dense phase pH. **(b)** FUS tagged with SNAP dense and dilute phase pH using PEG as phase separation trigger and fixed total protein concentration (2 µM), salt concentration (150 mM KCl) and buffer conditions (pH=7.3, 50 mM HEPES buffer). **(c)** FUS dense and dilute phase pH using high ionic strength as a phase separation trigger and fixed total protein concentration (2 µM), salt concentration (150 mM KCl) and buffer conditions (pH=7.3, 50 mM HEPES buffer). **(d)** FUS dense and dilute phase pH using low ionic strength as a phase separation trigger and fixed total protein concentration (2 µM) and buffer conditions (pH=7.3, 5 mM HEPES buffer).

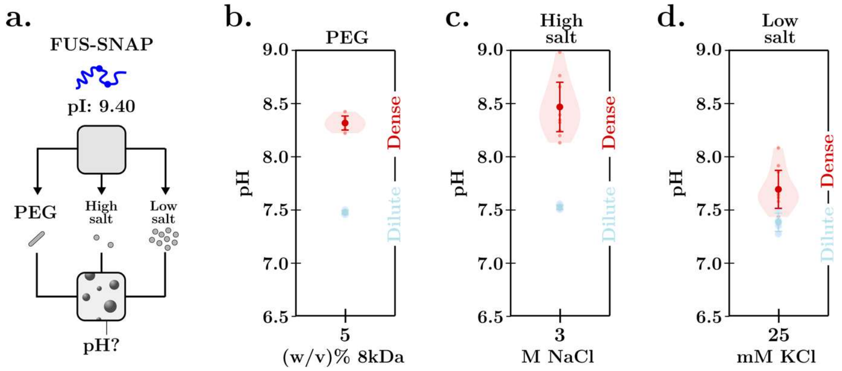


**Supplementary Figure 11: FUS phase separation under varying TRIS buffer strength.** Widefield imaging of FUS-EGFP condensates under varying concentrations of TRIS buffer at constant pH = 7.3 as well as constant total protein concentration (1 µM) and salt conditions (150 mM), highlighting that phase separation is observed at low TRIS concentrations.

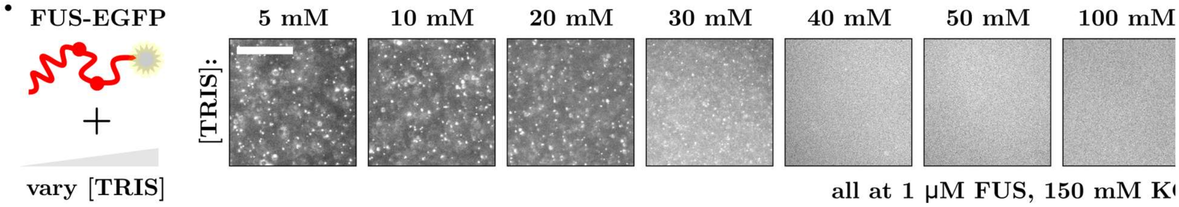

**Supplementary Figure 12: FUS dilute phase concentration at varying p(A)-RNA concentrations.** Dilute phase concentration readouts of FUS protein with varying p(A)-RNA concentrations. Phase separation is observed near composition at which the net charge of the pK + PGL3 mixture is predicted to be neutral at physiological pH (purple arrow).

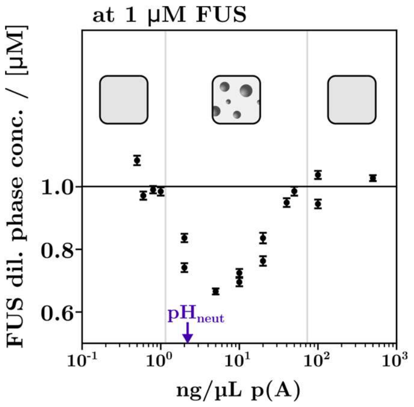


**Supplementary Figure 13: Changes in pI distribution considering folding architecture.** The plot displays the pI histogram of a subset of SwissProt/UniProt Human proteins using consideration of the folded state via application of the PypKA software[15] (blue) as well as from direct prediction of just the linearised sequence (orange).

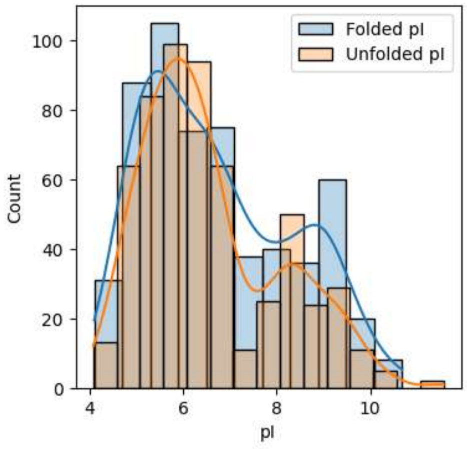


**Supplementary Figure 14: pI distribution of U2OS lysate proteome.** pI density distribution of all proteins detected in U2OS cell lysate both when considering only the individual proteins (grey) as well as their relative abundance (blue), data from [18].

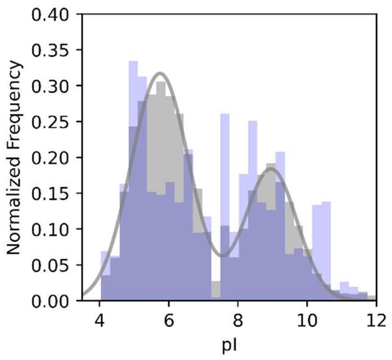

**Supplementary Figure 15: Overview of the side chain pKa's of the naturally charged amino acids.** Side chain pKa's of Aspartic acid, Glutamic acid, Histidine, Lysine and Arginine compared to the pI distribution of the full human proteome (grey background).

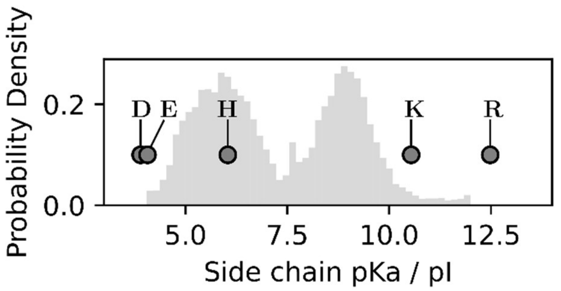


**Supplementary Figure 16: Proteome wide pI distribution for various species.** pI distribution of the full set of individual proteins contained in the proteome of the yeast species *Saccharomyces cerevisiae* (left, green), as well as the extremophiles *Acidithiobacillus thiooxidans* (middle, blue) and *Alkalihalobacillus clausii* (right, red) as obtained from Uniprot.

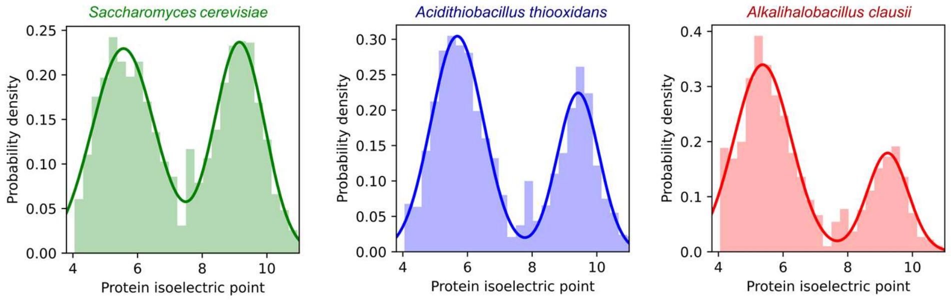


**Supplementary Figure 17: Comparison between the human and randomised proteomes.** **(a)** Distribution of isoelectric points of highly phase separation prone proteins (DeePhase Score > 0.8)[19] (top) as well as the full human proteome (bot). **(b)** Distribution of isoelectric points of randomised proteomes compared to the human proteome. (top) Human amino acid abundance probability and length distribution and (bot) fully randomised sequences.

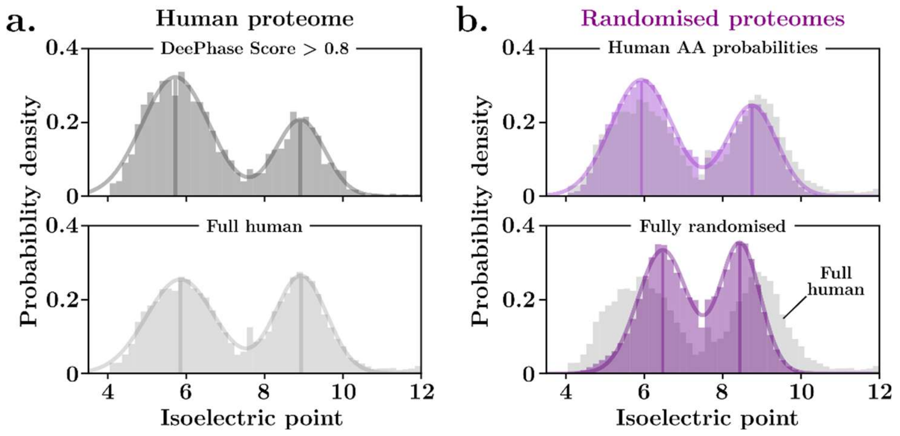

**Supplementary Figure 18: Electrochemical properties of an NMP1 lysate pulldown proteome. (a)** Distribution of isoelectric points of quantitative proteomics data obtained from NPM1 condensation in cellular lysate both considering only the individual proteins (top) and the relative abundance (bot). **(b)** Changes in the net charge of the NPM1 lysate pulldown proteome with varying pH as normalised by the total number of proteins, showing the comparison between considering just the individual proteins as well as the quantitative abundance data. The point at which the condensate proteome net charge is neutralised is further referred to as the condensate proteome mixture pI.

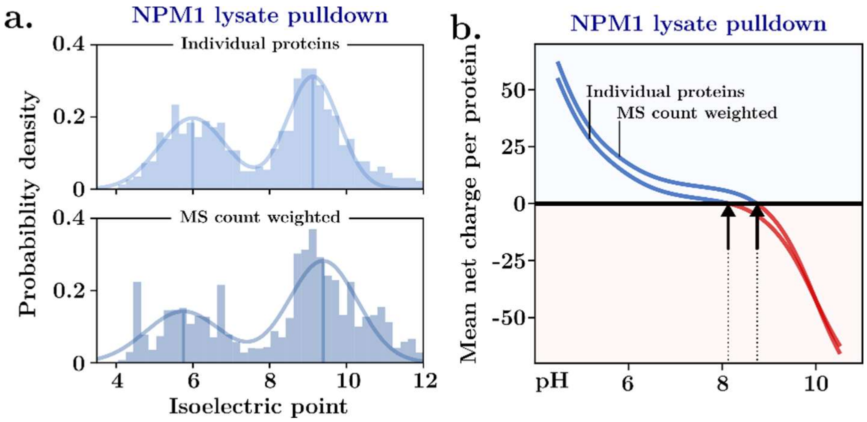

**Supplementary Figure 19: Property comparison of the Condensate Atlas.** **(a)** Schematic representation of the condensate Atlas and **(b)** box plot representation of the number of individual proteins per individual condensate network and percentage of phase separation prone proteins per individual condensate network in the Condensate Atlas. **(c)** Histogram distributions of characteristic features of the Condensate Atlas. Histograms of the number of individual proteins per condensate network, fraction of phase separation prone proteins, condensate cluster pI and the net charge change at the pI (from left to right) for all 85 entries. All histograms given in probability density. **(d)** Evaluation of the proteome pH response behaviour around the mixture pI. Change in the charge responsiveness to pH pertubations $-(\Delta C/\Delta pH)|_{pI}$ against condensate cluster pI. Grey box: highlighting the range of $-(\Delta C/\Delta pH)|_{pI}$ values of different condensate cluster with similar pI. **(e)** Schematic representation highlighting the diversity of potential pH gradients and pH buffering responses display by complex condensates.

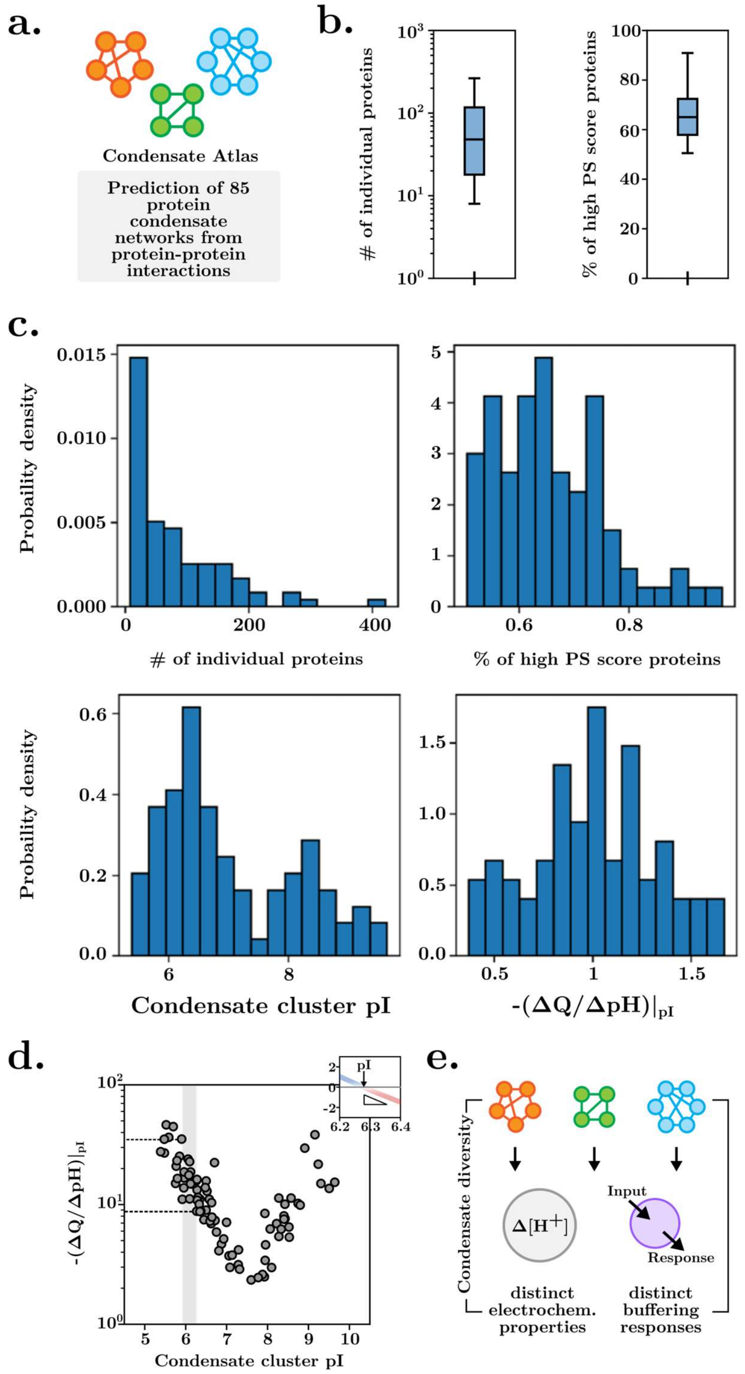
